# FMRP safeguards expression from long mRNAs by opposing repressive RNP states

**DOI:** 10.64898/2026.07.23.739932

**Authors:** Kaicheng Ma, Al Rohet Hossain, Kayla Judson, Thomas Cheng, Ethan J. Greenblatt

**Author notes:** Equal contribution (Kaicheng Ma, Al Rohet Hossain). Corresponding author: Dr. Ethan J. Greenblatt.

## Abstract

Fragile X messenger ribonucleoprotein (FMRP) is widely viewed as a translational repressor, yet its loss paradoxically compromises expression of exceptionally long, dosage-sensitive neurodevelopmental mRNAs. First shown in *Drosophila* oocytes, this selective loss extends to mouse cortex and hippocampus, but its mechanism remains unknown. Here, we uncover a protective function for FMRP: shielding long mRNAs from repression by Me31B/DDX6, a conserved RNA helicase that organizes repressive RNP complexes. In *Drosophila* oocytes, FMRP loss redirects target mRNAs into Me31B-marked P-bodies. Transcriptome-wide binding maps reveal extensive FMRP–Me31B co-occupancy, preferentially concentrated on long coding sequences whose translation depends on FMRP. Strikingly, disrupting Me31B-dependent repression preferentially enhances translation of FMRP targets in FMRP-deficient oocytes. We identify Rin/G3BP, Caprin, and Lig/UBAP2L as FMRP-associated cofactors supporting expression of distinct targets. Together, our findings challenge the prevailing view of FMRP as a translational repressor, revealing instead a mechanism that safeguards vulnerable neurodevelopmental mRNAs from inappropriate cytoplasmic silencing.

## INTRODUCTION

Neurodevelopment depends on the precise expression of many exceptionally large, dosage-sensitive genes.^1-3^ FMRP, encoded by *FMR1*, is a central post-transcriptional regulator of long mRNAs produced by this gene class.^4-8^ Loss of FMRP causes fragile X syndrome (FXS), a leading inherited cause of intellectual disability (ID) and autism spectrum disorder (ASD), and is also associated with fragile X-associated primary ovarian insufficiency.^9,10^ Dozens of FMRP-bound transcripts are intolerant to altered gene dosage, and are strongly associated with ID and ASD.^3,11,12^ FMRP targets contain coding regions three times longer than typical neuronal genes^8^ and encode proteins with diverse cellular functions, including chromatin regulators, cytoskeletal adaptors, ubiquitin ligases, and synaptic scaffolds.^11,13,14^ Despite its identification over 30 years ago, FMRP’s primary function and mode of action remain debated. This uncertainty stems in part from the difficulty of measuring RNA regulation *in vivo* in intact neurons, and from the many secondary consequences that follow the loss of a major gene regulator, which complicate phenotypic interpretation.^15^ The lack of disease-modifying therapies for FXS, despite extensive mechanism-based clinical efforts, underscores the need for a more complete understanding of FMRP’s primary molecular functions.^16,17^

Although FMRP has long been viewed primarily as a translational repressor,^11,18-22^ studies in *Drosophila* oocytes using ribosome profiling revealed that FMRP’s primary activity may instead be to promote protein synthesis. *Fmr1* knockdown oocytes showed ∼50% translational downregulation, specifically of large protein-coding mRNAs including many essential for neurodevelopment.^4^ Embryos derived from these oocytes exhibited reduced viability and nervous system defects, suggesting that FMRP loss results in disrupted development due to the underproduction of critical large neurodevelopmental proteins.^4^ The requirement for FMRP for efficient translation of long mRNAs was subsequently observed in the mouse hippocampus and cortex, where the most strongly FMRP-bound transcripts contained unusually long coding sequences and were also selectively reduced in protein output upon FMRP loss, as measured through ribosome profiling, TRAP-seq, CLIP-seq, proteomics, and capillary Western blot assays.^5,7,8,23^ FMRP’s conserved function is further supported by studies showing that expression of human FMRP rescues phenotypes in *Fmr1-*deficient *Drosophila*.^24^ Together, these findings establish *Drosophila* oocytes as a tractable system for dissecting the conserved yet unresolved mechanism of FMRP-dependent expression from long target mRNAs.

One possible mechanism is that FMRP protects long mRNAs from inappropriate entry into repressive RNP assemblies. Cytoplasmic mRNAs are packaged into dynamic ribonucleoprotein (RNP) complexes and can partition into higher-order RNP states that differ in molecular composition, translational activity, and downstream fate.^25-27^ P-bodies are enriched for conserved translational silencing machinery, including the DEAD-box helicase Me31B, the *Drosophila* ortholog of DDX6; the eIF4E-binding protein Cup/4E-T; and Tral/LSM14A/B.^28-33^ Although P-body accumulation is often associated with translational repression, visible P-bodies can form downstream of silencing and are not necessarily required for repression itself.^34^ Thus, P-body accumulation is a cytological marker of Me31B/DDX6-associated repressive RNP states, rather than direct evidence that P-bodies are the causal site of silencing. In oocytes and neurons, P-body-associated factors and related RNP assemblies regulate the storage, localization, stability, and translational repression of mRNAs, in some cases over extended periods.^35-40^ Stress granules and related assemblies form as a downstream consequence of integrated stress response-mediated global translation inhibition and contain distinct sets of RNA-binding proteins and translation factors, yet the physiological functions of stress granule machinery *in vivo* remain largely unresolved.^41-45^ Consistent with the idea that transcript length influences RNP fate, long transcripts are enriched in multiple RNA granule classes, and repressive Me31B-containing complexes can cooperatively associate with mRNAs in a length-dependent manner largely independent of sequence.^46-49^ Given this potential vulnerability of long mRNAs towards repression, we hypothesized that FMRP may not simply be a translational repressor or activator in isolation, instead, helping to control the balance between translation-competent and translationally repressed RNP states which are vital to the plasticity of neurons and germ cells.

Here, we test this idea in *Drosophila* oocytes, where maternal mRNAs are extensively partitioned between translation-competent and repressive cytoplasmic RNP states, and where maternal mRNAs guide subsequent embryonic neurodevelopment. Our findings reveal that FMRP limits the excessive silencing of a small subset of target mRNAs which are highly-bound by Me31B and depend on FMRP for their translation. This protective activity is supported by an FMRP-associated regulatory module of stress granule-linked proteins that promotes expression of distinct target subsets. Furthermore, we provide evidence that this activity is deeply conserved through comparisons with mammalian brain data. Together, these results define FMRP RNP complexes as safeguards against promiscuous repressive complexes, challenging the prevailing model of FMRP as a repressor and revealing a key mechanism by which FMRP loss disrupts neurodevelopmental gene expression.

## RESULTS

### FMRP limits long target mRNAs from accumulating in Me31B-marked P-bodies

To test how FMRP loss impacts target repression, we visualized endogenous FMRP target mRNAs in held mature oocytes naturally containing numerous P-bodies marked by Me31B-GFP, using an optimized single-molecule fluorescence *in situ* hybridization (smFISH) protocol (see Methods). We examined five previously identified FMRP target mRNAs, *poe*, *ctrip*, *ana3*, *HUWE1*, and *Nf1*, all of which encode large proteins whose mammalian orthologs are implicated in neurodevelopmental disease pathways and whose translation decreases by ∼2-fold in *Fmr1*-deficient oocytes.^4,8,11,50-53^ As a control for global changes in mRNA distribution, we also labeled bulk polyadenylated poly(A) mRNA. In control oocytes, FMRP target transcripts were broadly distributed throughout the cytoplasm and were detected both inside and outside Me31B-marked P-bodies (Fig. 1A-E). In contrast, FMRP depletion caused a substantial relocalization of each tested target mRNA in Me31B-marked P-bodies, with increases ranging from approximately two-fold to eight-fold depending on the transcript (Fig. 1A-E). Bulk polyadenylated RNA did not show a comparable redistribution (Fig. 1F), indicating that FMRP loss does not globally redirect maternal mRNAs into P-bodies. These data show that FMRP selectively limits P-body association of its long target mRNAs. Because P-body accumulation can reflect upstream RNP remodeling, we interpret this redistribution as evidence that FMRP loss shifts target mRNAs toward a repressive RNP state.

**Figure 1.**
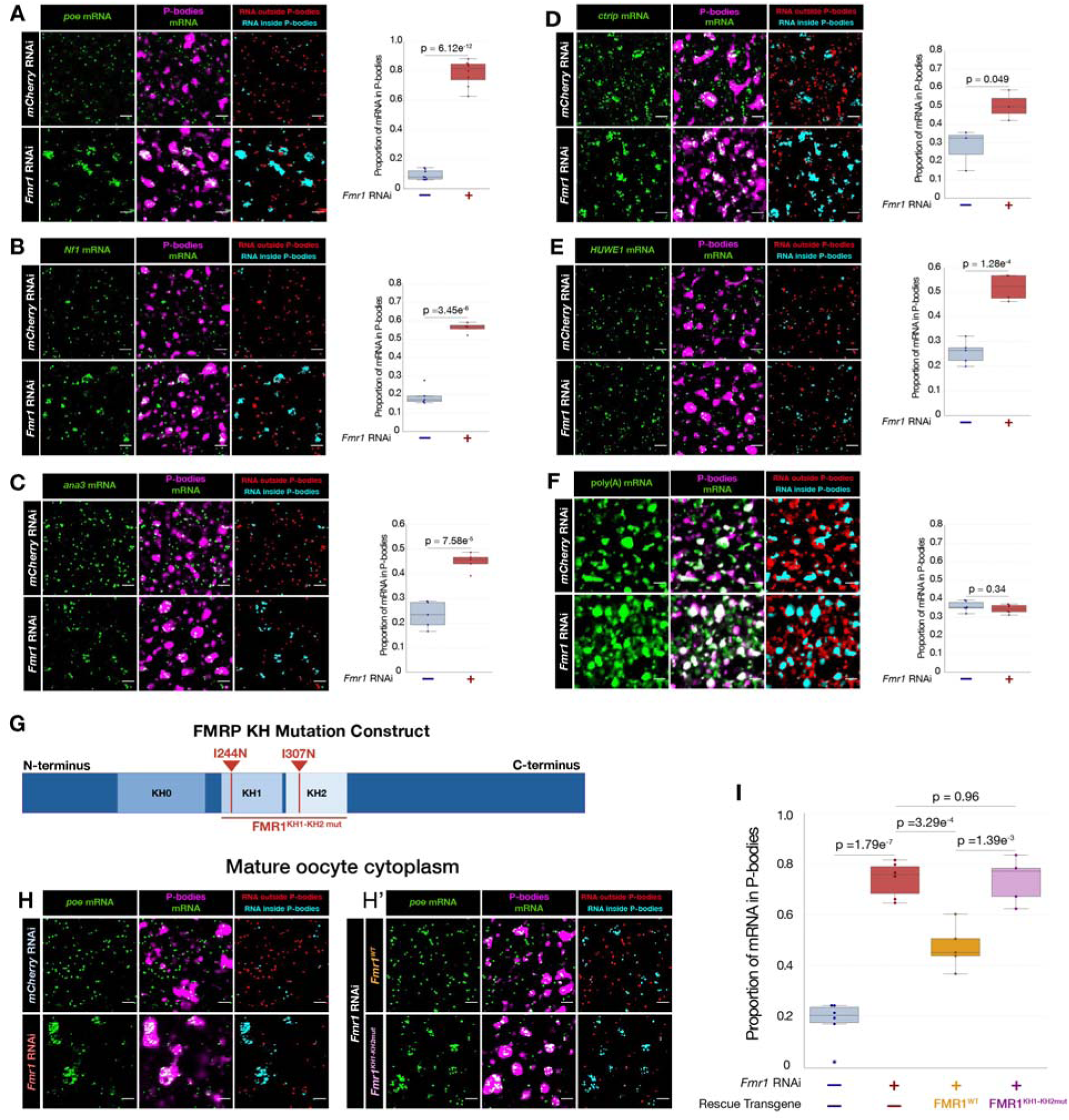
FMRP prevents long target mRNAs from accumulating in Me31B-marked P-bodies. **(A–F)** Representative single-molecule FISH (smFISH) images *(left)* and quantification *(right)* of *mCherry* RNAi or *Fmr1* RNAi mature oocytes showing increased P-body localization of FMRP target mRNAs *poe*, *HUWE1*, *Nf1*, *ctrip*, *ana3*, but not bulk poly(A) RNA. RNA is shown in green, and Me31B-GFP-labeled P-bodies are shown in magenta. Segmented smFISH signals outside or within P-bodies are shown in red and cyan, respectively. **(G)** Schematic of the FMRP KH domains. KH domains are indicated in dark blue, and arrows mark the I244N and I307N substitutions in *Fmr1*^KH1-KH2mut^. **(H and H□)** Representative Poe smFISH images and quantification **(I)** of control or *Fmr1* RNAi oocytes with or without expression of RNAi-resistant *Fmr1*^WT^ or *Fmr1*^KH1-KH2mut^ transgenes, showing partial rescue by wild-type *Fmr1* but not the KH domain double-mutant. Scale bars: 3 μm.

Although FXS is most commonly caused by a pathogenic CGG-repeat expansion that leads to *FMR1* transcriptional silencing, rare *FMR1* missense variants affecting FMRP’s KH RNA-binding domains can also cause FXS-like disease.^54-56^ We found that FMRP’s KH domains were required for FMRP-mediated exclusion of target mRNAs from P-bodies. *poe* mRNA localization outside of P-bodies was partially restored by re-expression of a wild-type *Fmr1* rescue transgene in *Fmr1* knockdown oocytes. However, expression of an FMRP mutant construct containing point mutations in two critical RNA-binding domains, KH1 and KH2, had no apparent rescue effect (Fig. 1G-I). These data suggest that pathogenic disruption of FMRP RNA-binding activity may promote inappropriate target mRNA association with P-body-associated repressive RNP states.

### FMRP and Me31B preferentially bind long coding sequences

The selective accumulation of FMRP targets in P-bodies raised the possibility that these transcripts are intrinsically vulnerable to engagement by Me31B-associated repressive machinery. To define the binding landscapes of FMRP and Me31B, we generated endogenously 3xFLAG-tagged FMRP and Me31B *Drosophila* lines using CRISPR and adapted improved individual-nucleotide resolution crosslinking and immunoprecipitation followed by sequencing (iiCLIP-seq) for mature oocytes.^57^ We optimized the iiCLIP protocol using a smaller anti-FLAG Fab, which essentially eliminated background RNA binding as compared to a full-length FLAG M2 antibody and enabled high-confidence library generation (Fig. 2A and Fig. S1). This approach yielded highly reproducible iiCLIP datasets for both FMRP and Me31B, with strong concordance between biological replicates (*R*² = 0.971 and *R*² = 0.970, respectively; Fig. 2B and 2C). Consistent with prior studies,^11^ FMRP peaks mapped primarily to coding sequences (CDS), which accounted for 57% of peaks (Fig. 2D). Me31B showed similar binding preference, with 56% of peaks mapping to the CDS (Fig. 2D). A small subset of mRNAs showed especially strong FMRP or Me31B binding, with iiCLIP scores substantially above the transcriptome average; these high-binding targets represented 5.7% and 6.1% of mRNAs, respectively (Fig. 2E and 2F). Surprisingly, across the full range of transcript binding levels, FMRP and Me31B occupancy were strongly correlated (*R*² = 0.863; Fig. 2G). FMRP and Me31B iiCLIP signal was also readily detected together across the coding regions of the long targets analyzed by smFISH (Fig. 2H).

**Figure 2.**
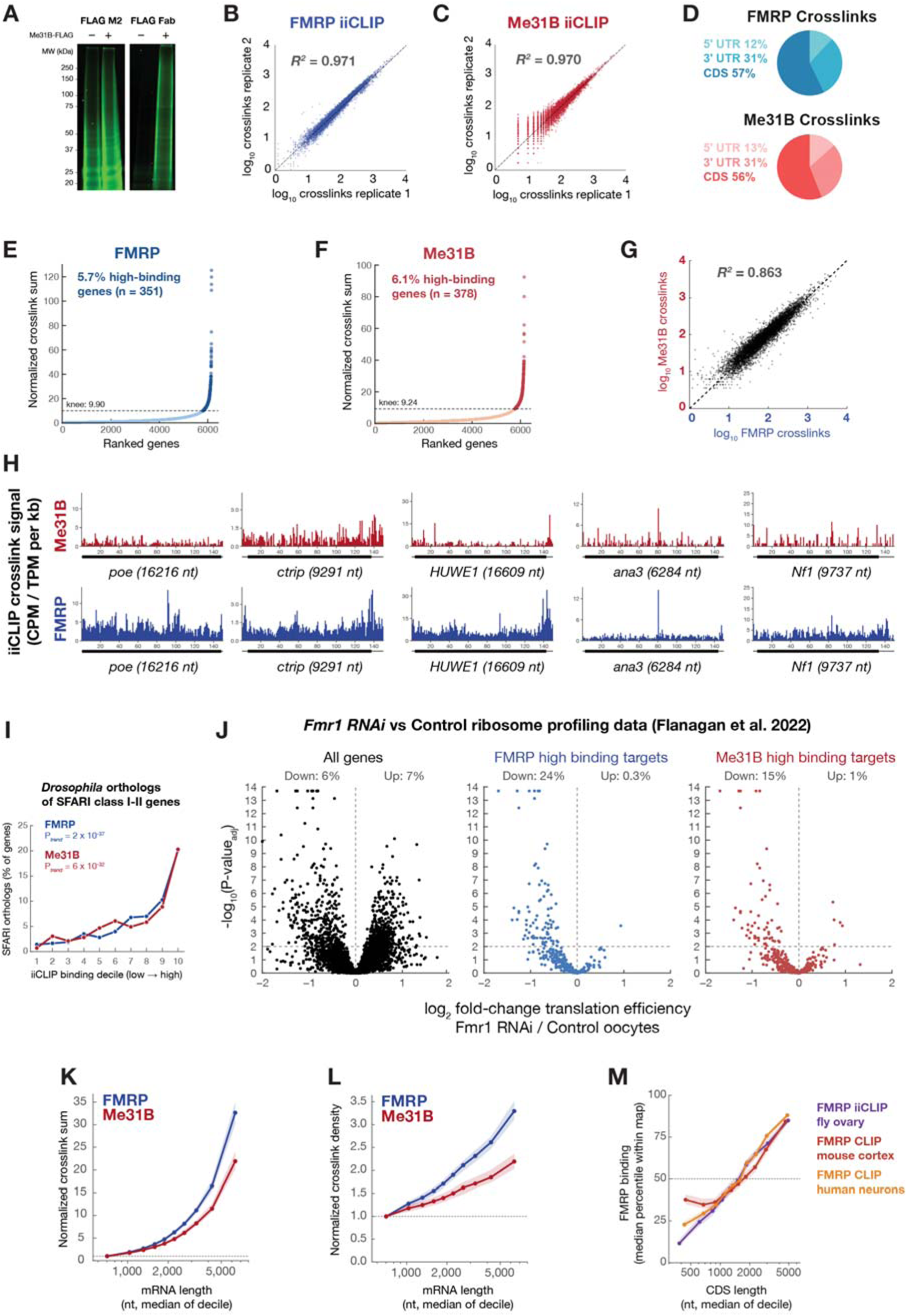
FMRP and Me31B co-occupy long coding sequences and define a repression-prone FMRP target set. (**A**) Comparison of background vs. specific RNA signal obtained from Me31B-FLAG co-immunoprecipitation of crosslinked RNA using FLAG M2 vs. Fab antibodies. (**B** and **C**) High reproducibility of FMRP-FLAG (**B**) and Me31B-FLAG (**C**) iiCLIP-seq crosslink values between biological replicates. (**D**) Distribution of exonic FMRP-FLAG (top) and Me31B-FLAG (bottom) crosslinks among the 5□UTR, CDS, and 3□UTR of bound mRNAs, showing majority of crosslinks are to the CDS. (**E** and **F**) mRNAs ranked by FMRP (**E)** or Me31B (**F**) crosslinks normalized to RNA abundance. The dashed line marks the knee of each curve which separates the high-binding targets (dark points) from all bound mRNAs. (**G**) Plot showing high transcript-level correlation between FMRP vs. Me31B iiCLIP-seq crosslink values. (**H**) FMRP (blue) and Me31B (red) crosslink profiles along five long mRNAs examined by smFISH in Figure 1. In the gene models below, thick bars are CDS and thin lines are UTRs. (**I**) *Drosophila* FMRP and Me31B crosslinks are highly enriched for *Drosophila* orthologs of human SFARI class I-II autism spectrum disorder-associated genes. *p*_trend_: Cochran–Armitage test for trend across binding deciles. (**J**) Volcano plots of the change in translation efficiency (TE) in *Fmr1* RNAi versus control oocytes.^8^ Panels show all analyzed mRNAs (left), FMRP high-binding targets (middle) and Me31B high-binding targets (right). (**K** and **L**) FMRP (blue) and Me31B (red) total (**K**) and length-normalized (**L**) binding increases with mRNA length. The median of RNA abundance normalized crosslinks is shown for each group, shading represents the 95% confidence intervals. (**M**) Increased FMRP binding density for long CDS mRNAs is conserved across species and tissues: *Drosophila* ovary FMRP iiCLIPseq (this study); mouse cortex FMRP HITS-CLIP^93^; human forebrain neuron *FMR1* CLIP^6^. Within each map, mRNAs were ranked by binding (percentile 0– 100). Shading represents the 95% confidence interval of the median.

To test whether FMRP and Me31B bind preferentially to the *Drosophila* orthologs of mRNAs linked to neurodevelopmental disease, we examined fly orthologs of SFARI class I–II autism risk genes (254 of 4,286 expressed mRNAs with a clear human ortholog). For both proteins, the fraction of SFARI orthologs rose with binding, from 1.4% (FMRP) and 0.7% (Me31B) in the lowest binding decile to 20.0% and 20.3% in the highest (*p*_trend_ = 2 × 10^−37^ and 6 × 10^−32^, Cochran-Armitage trend test; Fig. 2I). In all, 69 SFARI orthologs were high-binding targets of either FMRP or Me31B, and 50 of these were shared by both proteins (Table S1). Next, we integrated the iiCLIP datasets with ribosome profiling data from control and FMRP-depleted oocytes. Across the transcriptome, FMRP loss caused relatively modest changes in translational efficiency, with only a small fraction of transcripts significantly downregulated or upregulated (6% downregulated versus 7% upregulated; Fig. 2J). In contrast, FMRP high-binding targets were selectively sensitive to FMRP depletion and were exclusively reduced in translation: 24% were significantly translationally downregulated (adjusted *p* < 0.01), whereas 0.3% were significantly upregulated (Fig. 2J). Me31B high-binding targets showed a similar, though slightly less pronounced, bias toward reduced translation upon FMRP depletion (15% significantly downregulated versus 1% significantly upregulated; Fig. 2J).

Finally, we asked whether FMRP and Me31B binding scales with transcript length. FMRP binding increased strongly with mRNA length, both when measured as total RNA-abundance-normalized iiCLIP signal and when additionally normalized for transcript length (Fig. 2K, 2L, and Fig. S2). Long mRNAs therefore did not simply accumulate more FMRP signal because they provide more sequence space; they also exhibited higher FMRP binding density. Me31B showed a similar length-dependent pattern, with long transcripts accumulating both higher total Me31B signal and higher binding density than short transcripts (Fig. 2K and 2L). Upon comparing our FMRP iiCLIP analysis with published data, we observed that FMRP’s strong preference for long coding sequences is highly conserved across species (Fig. 2M). In all of the analyzed datasets (fly ovary iiCLIP, mouse cortex and human neuron CLIP), the median binding percentile ranking rose steadily with CDS length. Together, these data indicate that FMRP high-binding targets are long, strongly occupied by Me31B, and especially likely to lose translation when FMRP is depleted, supporting a model in which FMRP protects a repression-prone subset of Me31B-associated mRNAs.

### FMRP binding has a positive effect on translation in *Drosophila* ovaries and mouse cortex but no apparent effect in either direction in cultured cells

In contrast to our findings, FMRP has recently been proposed to repress translation through eIF4E-binding in cultured mammalian cells.^22,58^ We therefore compared the effects of FMRP binding on translation in *Drosophila* ovaries and the mouse cortex as compared to cultured cells, where FMRP was proposed to sequester its targets into repressive mRNPs, to determine whether FMRP binding has context-specific consequences which could explain the discrepancy (Fig. 3). In both the ovary and cortex datasets, we found a remarkably similar pattern, where FMRP binding was strongly predictive of diminished but not increased translation due to *Fmr1* deficiency in both of the *in vivo* datasets (Fig. 3A and 3B). FMRP mRNA CLIP targets in the highest bound classes were strongly biased towards diminished and not increased translation upon FMRP depletion in both datasets and to similar magnitudes, indicating that FMRP’s positive function is conserved across evolutionary distant organisms and tissues.

**Figure 3.**
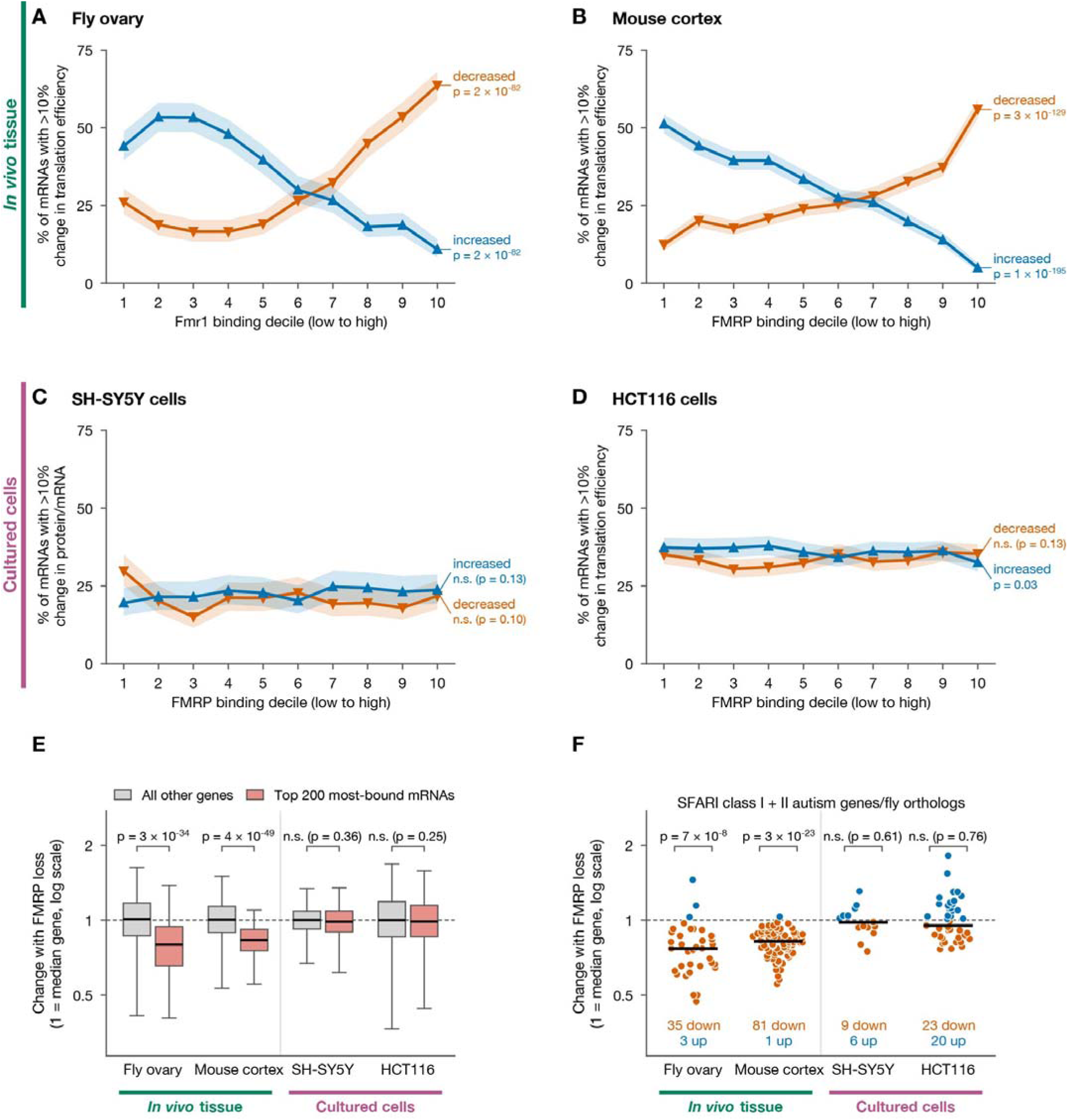
Translation of FMRP-bound mRNAs decreases after FMRP loss in fly ovary and mouse cortex but is largely unaffected in cultured cells. **(A–D)** FMRP binding is associated with reduced translation in fly ovaries and the mouse cortex but neither increased nor decreased translation in cultured cells. mRNAs binned into FMRP binding deciles based on CLIP-seq analysis (see Methods) (1, least bound; 10, most bound). Ribosome profiling translation efficiency **(A, B, D)** or protein/mRNA ratio **(C)** decreased (orange) or increased (blue) by more than 10% relative to the median gene after FMRP loss. Bands, Wilson 95% confidence intervals; *p* values, two-sided Cochran–Armitage test for trend. (A) *Drosophila* ovary, *Fmr1* RNAi. **(B)** Mouse cortex, *Fmr1* KO. **(C)** SH-SY5Y cells, *FMR1* KO. **(D)** HCT116 cells, *FMR1* KO. **(E)** Box plots showing reduced target translation after FMRP loss for the 200 most FMRP-bound mRNAs (red) for ovary and mouse cortex but not cultured cells datasets as compared to all other mRNAs (grey). *p* values, two-sided Mann–Whitney U test. **(F)** FMRP-bound fly and mouse orthologs of SFARI class I and II are strongly reduced in translation in fly ovary and mouse cortex but not cultured cell datasets. *p* values, two-sided sign test. n.s., not significant.

For the two cultured datasets analyzed, however, there was no significant effect in either direction when comparing strong vs. weakly FMRP-bound transcripts (Fig. 3C and 3D). When we stratified transcripts into deciles based on FMRP binding, we found no consistent relationship between binding strength and increased or diminished translation and protein to RNA ratios. There was also no significant difference in translation efficiency or protein to RNA ratios when we compared the most strongly FMRP-bound 200 genes to all other genes, as opposed to the ovary and cortex datasets where binding strength was strongly associated with reduced translation (Fig. 3E). This discrepancy was especially striking for genes known associated with ASD and ID (Fig. 3F). Whereas orthologs of SFARI class I and II ASD-associated genes were strongly biased towards downregulation in the fly ovary and mouse cortex datasets (35/40 and 81/82 downregulated, *p* = 7 × 10^−8^ and *p* = 3 × 10^−23^ respectively), there was no significant bias in either direction in *Fmr1* deficient SH-SY5Y or HCT116 cells (9/15 and 23/43 downregulated, *p* = 0.61 and *p* = 0.76 respectively).

Together, these data indicate that FMRP promotes rather than represses expression across evolutionary divergent tissues, with strong consequences on ASD-associated gene expression, but has little effect on translation efficiency in cultured mammalian cells, in contradiction to its proposed repressive mechanism there.

### Inhibition of Me31B-dependent repression increases translation in FMRP-deficient oocytes

The accumulation of FMRP targets in P-bodies upon FMRP loss could indicate that Me31B-dependent repression causes translational failure. However, P-body localization can also occur as a downstream consequence of reduced translation.^34^ To distinguish these possibilities, we performed a genetic interaction experiment that inhibited Me31B function in otherwise control or FMRP-deficient oocytes.

Prior work showed that a Me31B E208A substitution in the DEAD-box motif acts as a dominant-negative allele in the *Drosophila* ovary, leading bound transcripts to become translationally de-repressed.^59^ Because Me31B is essential for germline development, we found that low-level expression of a dominant-negative *me31B* transgene provided a way to disrupt Me31B-dependent repressive complex assembly while allowing oocyte development to proceed to completion. We reasoned that if FMRP protects its targets from Me31B-dependent repression, then Me31B inhibition through expression of Me31B^DN^ should have limited effects on FMRP targets when FMRP is present but should strongly de-repress these targets when FMRP is depleted. We therefore generated a GFP-tagged full-length *me31B* transgene carrying the dominant-negative E208A mutation (Me31B^DN^), expressed under UAS/GAL4 control to drive Me31B^DN^ specifically in the germline. Western blotting of oocyte extracts confirmed expression of the dominant-negative *me31B*^DN^*-GFP* transgene at 10-15% of the endogenous Me31B protein level, and expression was similar with or without *Fmr1* RNAi (Fig. 4A). We performed ribosome profiling and RNA-seq on control oocytes and oocytes expressing germline Me31B^DN^, with or without *Fmr1* RNAi. To control for global differences in RNA recovery and translation measurements, we added *Drosophila pseudoobscura* ovary extract spike-in to each sample.

**Figure 4.**
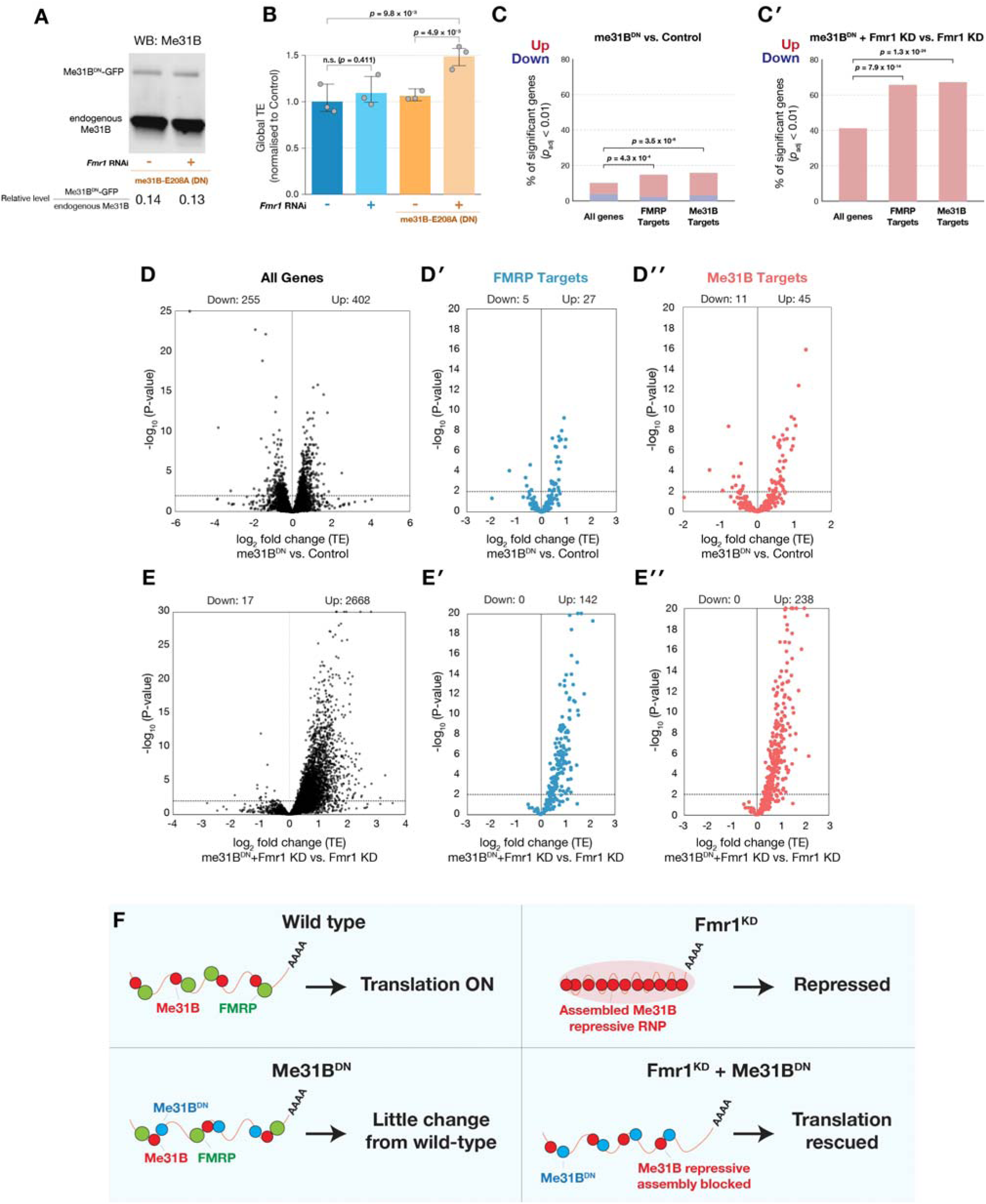
Expression of dominant-negative Me31B restores translation in FMRP-deficient oocytes. **(A)** Immunoblot of whole ovary for Me31B, showing that the ratio of dominant-negative E208A mutant Me31B^DN^-GFP vs. endogenous Me31B levels are similar in control vs. *Fmr1* RNAi oocytes. **(B)** Plots showing spike-in normalized global translation efficiency (TE) values across genotypes: control oocytes vs. *Fmr1* RNAi oocytes with or without additional expression of *me31B*^DN^. Only the combined condition (*Fmr1* RNAi plus Me31B^DN^) significantly altered global TE, increasing it 1.5-fold, relative to control oocytes. Bars show the mean of three biological replicates; error bars, 95% bootstrap confidence intervals; statistical significance determined by two-sided permutation test. **(C)** Fractions of genes with significantly increased or decreased TE (adjusted *p* < 0.01) among all genes, FMRP targets, and Me31B targets. Comparisons show Me31B^DN^ vs. control **(C)** and Me31B^DN^ plus *Fmr1* RNAi vs. *Fmr1* RNAi alone **(C□)**. Enrichment relative to the corresponding non-target background was assessed using Fisher’s exact test and was significant only among genes with increased TE. Both FMRP and Me31B targets were significantly enriched among genes with increased TE in each comparison, with a modest enrichment in the Me31B^DN^ vs. control comparison and a substantially stronger enrichment in the Me31B^DN^ plus *Fmr1* RNAi vs. *Fmr1* RNAi alone comparison. Red indicates increased TE, and blue indicates decreased TE. **(D–D□)** Volcano plots of TE changes in Me31B^DN^ vs. control oocytes for **(D)** all analyzed genes, **(D□)** FMRP targets, and **(D□)** Me31B targets, showing a slight bias towards increased TE for all gene classes. **(E–E□)** Volcano plots showing TE changes in Me31B^DN^ expression plus *Fmr1* RNAi versus *Fmr1* KD alone for **(E)** all genes, **(E□)** FMRP targets, and **(E□)** Me31B targets, showing a large bias towards increased TE values for all gene sets. **(F)** Model for cooperative repression of FMRP targets. When FMRP is present, its long targets remain relatively protected from Me31B-mediated silencing, so Me31B^DN^ alone has limited effect on their translation. When FMRP is depleted, these targets become vulnerable to Me31B-dependent repressive complex assembly and lose translation. Disrupting Me31B-dependent repression with Me31B^DN^ in this sensitized background de-represses and restores translation of FMRP targets.

Interestingly, *Fmr1* RNAi or Me31B^DN^ expression alone did not significantly increase global translational efficiency (Fig. 4B). However, we observed a striking synthetic interaction when co-expressing these two constructs. Combining *Fmr1* RNAi with Me31B^DN^ caused a ∼50% increase in global translational efficiency relative to control and to Me31B^DN^ alone (both *p* < 0.01, Fig. 4B).

We next asked whether this synthetic de-repression was enriched among FMRP and Me31B high-binding targets. In the presence of FMRP, Me31B^DN^ caused modest changes across all expressed genes: 402 transcripts showed increased and 255 showed reduced translation efficiency relative to control oocytes (Fig. 4C, 4D). Both high-binding target sets were modestly but significantly biased toward translational upregulation, to a similar degree: 45 Me31B highly-bound transcripts increased and 11 decreased (12.7% upregulated, *p* = 3.5 × 10^−6^; Fig. 4D□), and 27 FMRP highly-bound targets increased and 5 decreased (12.5% upregulated, *p* = 4.3 × 10^−4^; Fig. 4D□), compared with 6.2% upregulated of all genes. Thus, when FMRP is intact, Me31B inhibition produces limited de-repression, and does so comparably for both target sets.

In contrast, Me31B^DN^ expression produced a much stronger effect in FMRP-depleted oocytes that extended beyond FMRP targets, increasing translational efficiency across a broad fraction of the transcriptome (Fig. 4C-E, Fig. S3). However, within this global response, FMRP high-binding targets were disproportionately affected. Of 216 high-binding targets, 142 (∼65%) increased in translational efficiency and none decreased, compared with ∼40% increased translation efficiency for all genes irrespective of FMRP binding status (*p* = 7.9 × 10^−14^; Fig. 4C□, 4E□). Me31B high-binding transcripts showed a similarly directional response, with 238 increasing (67%; *p* = 1.3 × 10^−24^; Fig. 4E□) and none decreasing. The broader synthetic response may partly reflect impaired RNA release by ATPase-defective Me31B^DN^, ^60^ potentially leading to more widespread RNA association. Given the strong correlation between FMRP and Me31B binding across transcripts, this could explain why the combined perturbation affects targets beyond those most strongly bound by the wild-type proteins.

Together, these genetic interaction data support a model in which translational failure of FMRP targets results from excessive Me31B-dependent repression. When FMRP is present, long targets remain relatively protected from Me31B-mediated silencing, so Me31B^DN^ has a limited positive effect on their translation. When FMRP is depleted, these targets become vulnerable to Me31B-dependent repressive complex assembly, accumulate in P-bodies, and lose translation. Disrupting Me31B-dependent repression in this sensitized background thus restores translation of FMRP targets (model Fig. 4F).

### FMRP promotes formation of translation-associated Poe/UBR4 particles

FMRP has been associated with cytoplasmic RNP granules in neurons and oocytes, including structures containing markers of P-bodies and stress granules, and is associated with positive sites of translation at neuronal mitochondria.^35,61-63^ In contrast, liquid-liquid phase separation of the FMRP paralog FXR1 is associated with positive translational activity.^64^ Because P-bodies and stress granules are often associated with translational repression or stalled translation, we next asked whether FMRP-associated RNP particles in the oocyte are uniformly repressive or whether some instead mark translation-competent states.

We focused on *poe*/*UBR4*, a long FMRP target whose translation is reduced ∼50% upon FMRP depletion and whose protein product forms large spherical particles in developing egg chambers.^4^ In late stage 10 egg chamber nurse cells, the germline cells which produce the cytoplasm subsequently transferred to the oocyte, both *poe* mRNA and Poe protein formed prominent cytoplasmic particles at comparable densities (Fig. 5A). These particles, each containing an average of ∼55 *poe* mRNA molecules (Fig. S4A), were eliminated in *poe* RNAi and *Fmr1*-null follicles, confirming that the punctate smFISH signal represented genuine multi-copy *poe* mRNA assemblies rather than nonspecific probe accumulation and that their formation depended on the presence of FMRP (Fig. 5B). Poe particles were formed transiently during follicle development and were largely absent from completely mature stage 14 oocytes (Fig. 1A). By contrast, *Dhc64C* mRNA, another transcript that formed cytoplasmic particles in late follicles whose translation was unaffected by the loss of *Fmr1*, showed no change in either *Fmr1-*null or *poe* RNAi follicles (Fig. S4A and S4B), indicating that the loss of Poe particles is specific rather than a general disruption of cytoplasmic mRNA granules in *Fmr1*-deficient cells.

**Figure 5.**
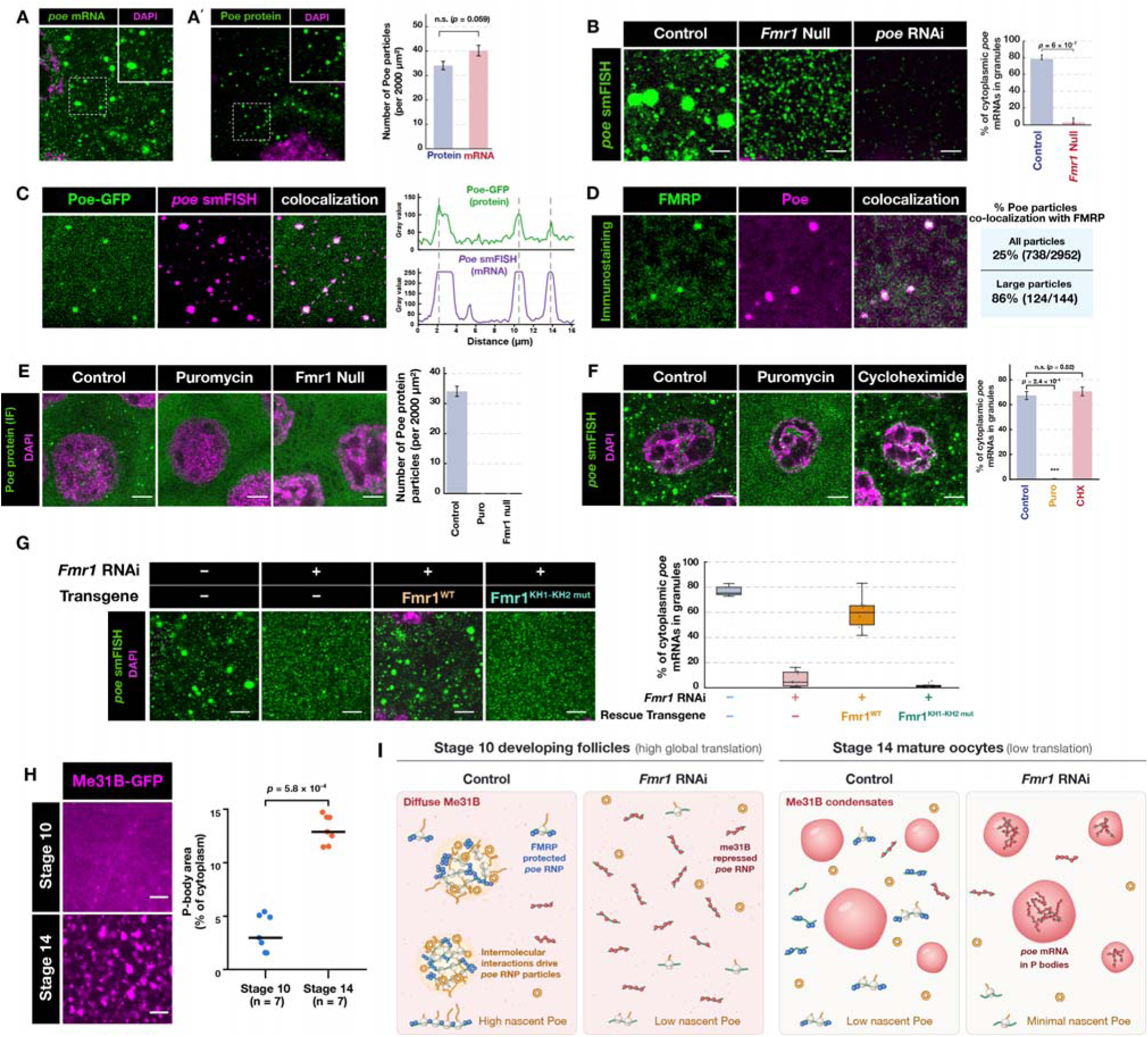
FMRP promotes formation of translation-associated Poe RNP particles in late-developing stage 10 follicles. **(A, A□)** *poe* smFISH (A) and Poe immunostaining (A□) of stage 10 nurse cells, showing similar patterns of mRNA and protein particles in separately analyzed follicles; insets show magnified regions. **(A□)** Quantification showing comparable numbers of Poe protein and *poe* mRNA particles. **(B)** *poe* smFISH in control, *Fmr1*-null, and *poe* RNAi stage 10 nurse cells, showing that FMRP is required for formation of cytoplasmic *poe* mRNA granules. **(C)** Full-length Poe-GFP protein and *poe* smFISH signals co-localize in cytoplasmic particles; line scan at right shows coincident protein and mRNA peaks. **(D)** Immunostaining of FMRP and Poe protein showing co-localization within Poe particles. **(E)** Poe protein immunostaining of control, puromycin-treated, and *Fmr1*-null stage 10 nurse cells, showing loss of Poe protein particles after polysome disassembly by puromycin treatment and in *Fmr1*-null cells. **(F)** *poe* smFISH of control, puromycin-treated, and cycloheximide (CHX)-treated stage 10 nurse cells, showing loss of *poe* mRNA granules with puromycin but not CHX treatment, which preserves polysomes. **(G)** *poe* smFISH showing partial rescue of Poe particle formation by a full-length *Fmr1*^WT^ transgene but not by an *Fmr1*^KH1-KH2mut^ transgene. **(H)** Me31B-GFP in stage 10 nurse cells and stage 14 mature oocytes, showing dispersed Me31B signal in stage 10 and sharply increased P-body formation at stage 14; quantification at right. **(I)** Model for *poe* mRNA dispersal in stage 10 nurse cells and P-body condensation in stage 14 mature oocytes. FMRP protection promotes high levels of Poe protein synthesis, which drives particle formation through intermolecular interactions between nascent Poe dimers and FMRP multimers. Loss of FMRP leads to reduced nascent Poe, disrupting particle formation and redistributing *poe* RNA into dispersed Me31B RNPs. In stage 14, reduced global translation levels leads to a loss of translation-associated Poe particles, and *Fmr1* knockdown leads to *poe*P-body accumulation via Me31B.

To determine whether *poe* mRNA and Poe protein occupy the same structures, we generated animals expressing endogenously tagged Poe-GFP and combined Poe-GFP imaging with *poe* smFISH. Because GFP was fused to the C-terminus of Poe, this reporter marks full-length Poe protein rather than an N-terminal fragment or stalled translation intermediate. *poe* mRNA granules colocalized with Poe-GFP protein particles, and line-scan analysis confirmed coincident mRNA and protein enrichment within individual particles (Fig. 5C). This pattern was not a general property of visible mRNA particles: *Dhc64C* mRNA also did not colocalize with Poe-GFP (Fig. S4C).

We next asked whether Poe particles are associated with FMRP. Immunostaining showed FMRP colocalizing with Poe protein particles, particularly with larger particles: 86% of large Poe particles greater than 0.2 μm^2^ contained detectable FMRP signal, compared with 25% of all Poe particles regardless of size (Fig. 5D). This size dependence may reflect the limited sensitivity of detecting particle-enriched FMRP above the strong diffuse cytoplasmic FMRP signal, rather than an exclusive association of FMRP with large Poe particles.

We next tested whether Poe particles depend on ongoing translation and FMRP. Puromycin, which releases nascent chains and disassembles polysomes, rapidly dispersed Poe-GFP particles following a 15-minute treatment. Poe-GFP particles were also lost in *Fmr1*-null follicles (Fig. 5E). Thus, full-length Poe protein particles depend on both active translation and FMRP. *poe* mRNA granules showed a similar translation-dependent behavior. Puromycin rapidly dispersed *poe* mRNA granules, whereas cycloheximide, which freezes elongating ribosomes on mRNAs, did not (Fig. 5F). This pharmacological sensitivity indicates that Poe particles are maintained by ongoing translation rather than by a stalled, translationally repressed RNP state.

Consistent with a requirement for FMRP’s RNA-regulatory activity, *poe* mRNA granule formation was restored by re-expression of wild-type FMRP in *Fmr1* RNAi follicles, but not by an *Fmr1* transgene carrying KH-domain mutations (Fig. 5G).

Stage 10 nurse cells and mature stage 14 oocytes differ sharply in their levels of translation as well as in how Me31B is organized, providing a potential explanation for why *poe* mRNA was dispersed following *Fmr1* knockdown in stage 10 nurse cells but concentrated into P-bodies in *Fmr1* knockdown mature oocytes. Me31B-GFP was largely diffuse in stage 10 nurse cells, whereas in stage 14 oocytes, which have much reduced levels of translation^65^, Me31B was mostly condensed into P-bodies occupying an 8-fold greater cross-sectional area within the cytoplasm (Fig. 5H; *p* = 5.8 × 10^−4^). These observations suggest a model (Fig. 5I). In wild-type stage 10 nurse cells, FMRP-bound *poe* polysomes assemble into translation-associated particles driven to coalesce by the known dimerization properties of both Poe protein^66^ as well as RNA-binding proteins such as FMRP which support high levels of nascent Poe synthesis and are also known to multimerize.^67^ Without FMRP, *poe* mRNA instead remains in dispersed, Me31B-repressed mRNPs and Poe synthesis falls. In stage 14 oocytes, where translation is globally low, FMRP loss shifts *poe* mRNA into a repressed state and *poe* mRNA is relocalized into P-bodies via Me31B.

Together, these data show that Poe particles contain *poe* mRNA and full-length Poe protein, require ongoing translation and FMRP KH-domain function, and are distinct from nonspecific cytoplasmic mRNA particles. These findings support a model in which FMRP promotes formation of translation-associated Poe particles, providing a visible readout of FMRP-dependent target expression.

### A Poe particle screen identifies conserved stress granule-linked FMRP-associated cofactors

The strong dependence of Poe particles on FMRP suggested that particle formation could provide a sensitive readout for factors required for FMRP-dependent target regulation. FMRP loss reduces *poe* translation by approximately two-fold,^4^ but nearly abolishes visible Poe particle formation. Having established that Poe particles contain both *poe* mRNA and full-length Poe protein, we used *poe* smFISH as a quantitative readout of Poe particle formation, reasoning that cofactors acting with FMRP to promote *poe* translation might be identified by screening for perturbations that similarly disrupt *poe* mRNA localization into large particles.

We performed an smFISH-based screen in stage 10 nurse cells, measuring the fraction of total *poe* mRNA signal localized to particles after depletion or mutation of candidate germline RNA-binding factors and ubiquitin-associated factors. Ubiquitin-associated candidates were included because Poe/UBR4 itself is a large ubiquitin ligase, raising the possibility that ubiquitin-linked pathways influence Poe particle assembly or stability. The screen included 151 RNAi or mutant lines targeting 117 genes, with *Fmr1* knockdown serving as a positive control and *mCherry* RNAi or a wild-type strain (Oregon-R) serving as matched controls. Representative images showed that most perturbations had little effect on *poe* mRNA localization, whereas *Fmr1* depletion and a subset of candidate perturbations strongly disrupted Poe particle formation (Fig. 6A). Quantification across the screen identified a small group of top-ranked hits, including *Fmr1*, *Rin/G3BP*, *CG32344/DDX54*, *Lig/UBAP2L*, *Otu/OTUD4*, and *Capr/CAPRIN1* (Fig. 6B). Interestingly, Rin, Lig, Otu, and Capr and their mammalian orthologs co-localize in stress granules together with FMRP.^68,69^ Conversely, depletion of several deubiquitinases increased *poe* mRNA localization to particles, suggesting that ubiquitin-regulated processes may tune the assembly or persistence of Poe particles (Fig. 6B).

**Figure 6.**
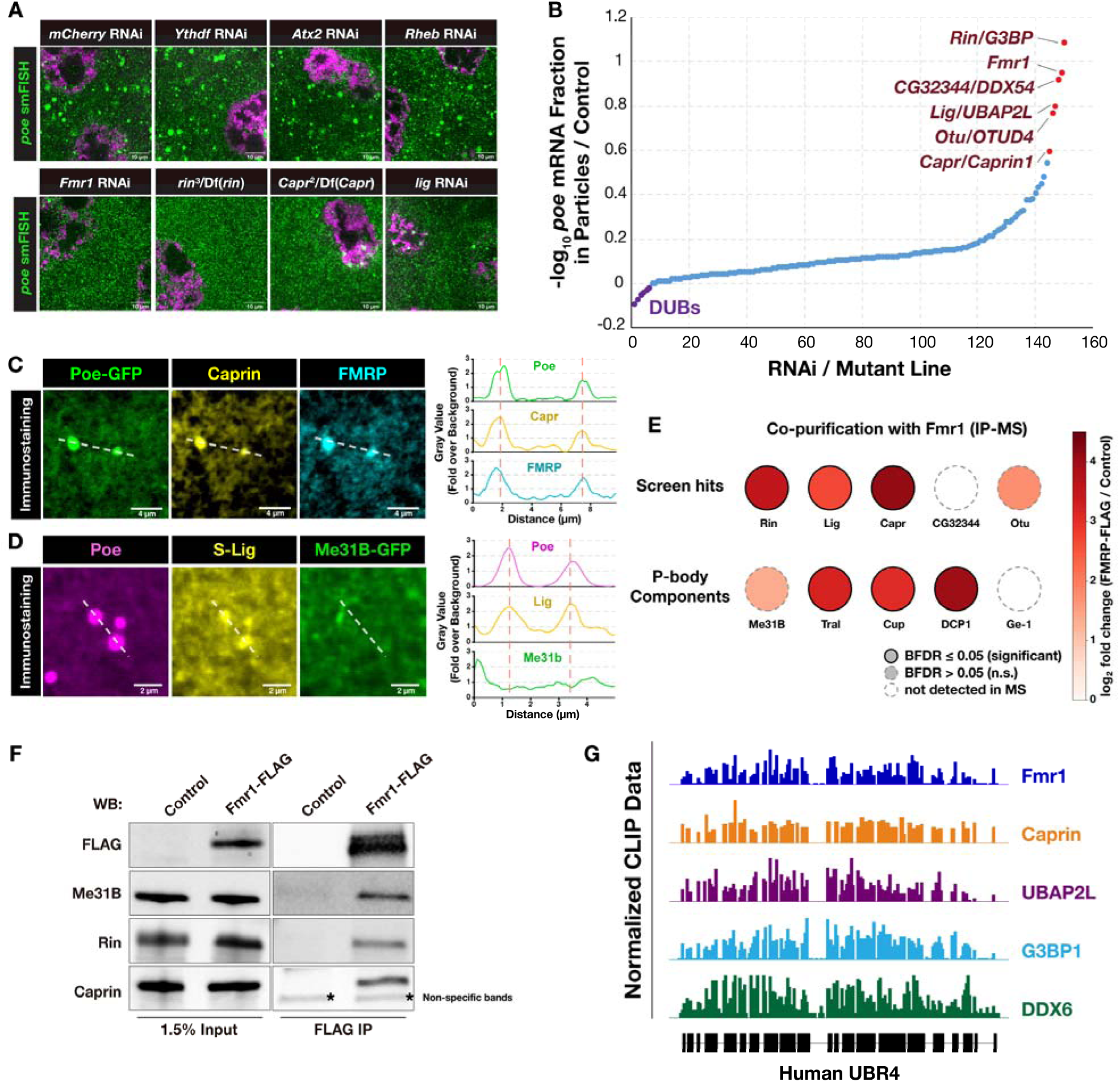
A candidate-based screen identifies stress granule-linked proteins Rin/G3BP, Capr/Caprin1, and Lig/UBAP2L as FMRP-associated cofactors. **(A)** Examples of non-hits (top row) and hits (bottom row) from a candidate-based *poe* smFISH screen for factors required for Poe particle formation. *poe* mRNA is shown in green and DAPI in magenta. **(B)** Ranked plot showing changes in *poe*mRNA particle enrichment across 151 mutant or RNAi lines targeting germline RNA-binding proteins and ubiquitin-associated factors, with top-ranked perturbations highlighted that disperse *poe* RNA particles (red) or increase *poe* RNA enrichment in particles (purple). **(C)** Immunostaining showing co-localization of Poe-GFP, Caprin, and FMRP within particles; line scans at right show coincident peaks. **(D)** Immunostaining of Poe, S-Lig, and Me31B-GFP showing co-localization of Poe and S-Lig but not Me31B; line scans are shown at right. **(E)** Dot plot of proteins co-purifying with FLAG-tagged FMRP by triplicate IP-MS, showing screen hits and P-body-associated components. Color indicates log_2_ fold change relative to control IP, solid outlines indicate significance (BFDR ≤ 0.05), dashed outlines indicate no significance (BFDR > 0.05) and empty fill indicates proteins not detected. Among the screen hits, Rin, Lig, and Capr are significantly enriched, whereas Otu does not meet significance and CG32344 is not detected. Among P-body-associated factors, FMRP co-purifies with Me31B, Tral, Cup, and DCP1, whereas the decapping-promoting protein Ge-1/EDC4 is not detected. **(F)** Western blots of FLAG-tagged FMRP complexes following co-immunoprecipitation, supporting association with Me31B, Rin, and Caprin. **(G)** Human iCLIP and eCLIP tracks over *UBR4*, the human ortholog of *Drosophila poe*, showing binding of FMRP, Caprin1, UBAP2L, G3BP1, and DDX6, the Me31B ortholog.

To determine which hits localize to Poe particles, we next examined candidate factors by imaging. Poe-GFP, Capr, and FMRP colocalized within the same cytoplasmic particles, and line-scan analysis confirmed coincident enrichment of all three signals (Fig. 6C). Tagged S-Lig also localized to Poe particles, whereas Me31B-GFP, which was mostly diffuse at this stage, was not similarly enriched at these sites (Fig. 6D). Thus, Poe particles contain a subset of stress granule-associated FMRP cofactors and are distinct from the Me31B-marked P-body state into which FMRP targets accumulate upon FMRP loss.

To distinguish candidate cofactors that physically associate with FMRP from factors that indirectly affect Poe particle formation, we performed proteomic analysis of FMRP immunoprecipitates. Mass spectrometry of FMRP-FLAG complexes identified Rin, Lig, and Capr as enriched FMRP-associated proteins, along with several P-body-associated factors, including Me31B, Tral, Cup, and DCP1 (Fig. 6E). In contrast, CG32344/DDX54 was not detected, and other screen hits were less strongly enriched. Notably, we did not recover Ge-1, the *Drosophila* ortholog of the decapping scaffold EDC4. FMRP thus associates with the storage and translational-repression module of P-bodies (Me31B, Tral, Cup) and the decapping activator DCP1, but not with the EDC4-type scaffold that couples decapping to 5□ to 3□ mRNA decay, indicating engagement of the repressive rather than the committed mRNA-degradation arm.^70^ Orthogonal FLAG immunoprecipitation followed by Western blotting confirmed recovery of Me31B, Rin, and Capr with FMRP-FLAG (Fig. 6F). These data indicate that FMRP associates with both translation-supportive granule factors and P-body repression machinery, consistent with a role in controlling transitions between alternative cytoplasmic RNP states.

Finally, we asked whether this FMRP-associated regulatory network is conserved on the mammalian *UBR4* transcript. Human iCLIP tracks from previous studies^71-75^ showed binding of FMRP, Caprin1, UBAP2L, G3BP1, and DDX6 across UBR4 (Fig. 6G). Thus, the Poe particle screen identified conserved FMRP-associated RNP factors that also engage the human UBR4 transcript and suggests that FMRP acts together with a translation-supportive RNP module to preserve target mRNA availability to gene expression machinery.

### FMRP-associated RNP factors act modularly to support target expression

We next asked whether FMRP-associated RNP factors support expression of FMRP targets beyond *poe* mRNA. Western blot analysis of established FMRP targets showed that FMRP depletion reduced steady-state levels of Poe, Ana3, and Vps13 proteins (Fig. 7A and S5A-C). Loss of Rin, Lig, or Capr produced target-specific effects. Poe protein was reduced most strongly after Rin or Capr loss, Ana3 protein was particularly sensitive to Lig depletion, and Vps13 protein was reduced to varying degrees across FMRP-associated cofactor perturbations. Yolk proteins, used as internal loading controls, were unaffected (Fig. 7A). These protein effects were not explained by corresponding decreases in target mRNA abundance. Quantification of Western blot signal together with RNA-seq showed that FMRP-associated cofactor perturbations reduced target protein levels while mRNA levels were either unaffected or increased in the corresponding transcripts (Fig. 7B). Global mRNA abundance, measured by spike-in-normalized RNA-seq, was unchanged across the *Fmr1*, Rin, Lig, and Capr perturbations (Fig. S5D), confirming that these effects do not reflect a change in total transcript levels. Thus, Rin, Lig, and Capr support expression of these FMRP targets primarily through post-transcriptional mechanisms.

**Figure 7.**
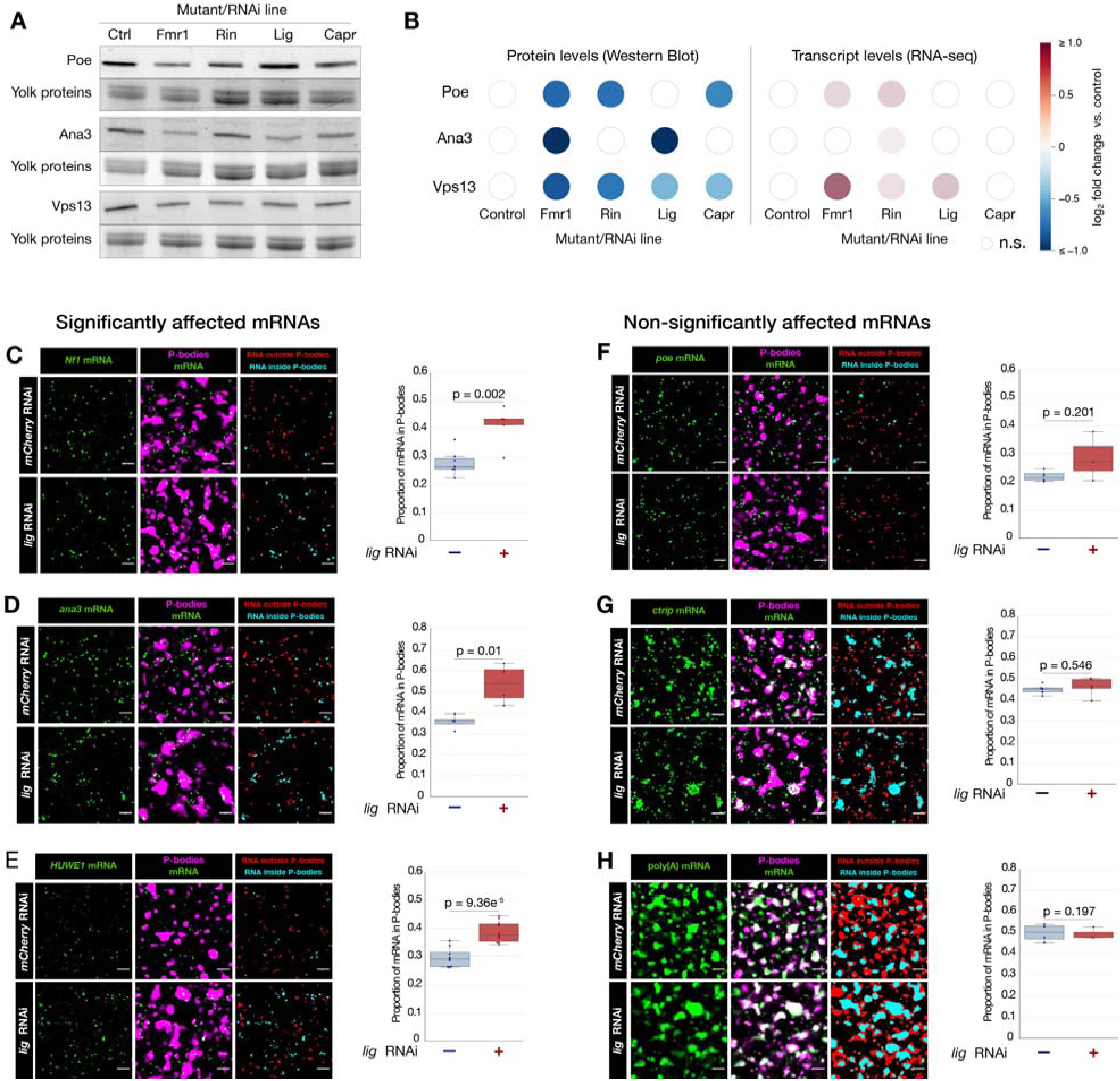
Rin/G3BP, Caprin, and Lig/UBAP2L are differentially required for FMRP target expression. **(A)** Western blot analysis showing differential reduction of Poe, Ana3, and Vps13 protein levels following loss of *Fmr1*, *rin*, *lig*, or *Capr*. **(B)** Quantification of protein abundances and corresponding transcript abundances measured by RNA-seq (right), showing that reductions in protein levels are not caused by reductions in mRNA. **(C–H)** Representative smFISH images and quantification of FMRP target mRNAs and poly(A) mRNA from control (*mCherry* RNAi) or *lig* RNAi mature oocytes expressing Me31B-GFP. RNA is shown in green, and Me31B-GFP-labeled P-bodies are shown in magenta. Masked RNA signal outside and within P-bodies is shown in red and cyan, respectively. Scale bars: 3 μm. **(C–E)** *Nf1*, *ana3*, and *HUWE1* mRNAs displayed significantly increased enrichment within P-bodies following *lig* depletion. **(F–H)** P-body enrichment of *poe* and *ctrip* or bulk poly(A) mRNAs was not significantly altered following *lig*depletion. Quantification is shown to the right of the corresponding representative images; *p* < 0.05 was considered statistically significant.

To determine whether this regulation includes protection from P-body sequestration, we examined whether individual FMRP-associated cofactors contribute to the anti-repressive activity revealed by FMRP loss. Rin^null^ oocytes showed a pronounced disruption of Me31B-GFP-marked P-bodies (Fig. S6), which became markedly smaller and less prominent, confounding direct measurement of target partitioning into P-bodies. In contrast, Lig depletion left Me31B-GFP-marked P-bodies largely intact, allowing target mRNA partitioning to be quantified.

We therefore co-imaged endogenous FMRP target mRNAs by smFISH together with Me31B-GFP-marked P-bodies in mature oocytes after Lig depletion (Fig. 7C-H). *Nf1*, *ana3*, and *HUWE1* mRNAs each showed significantly increased P-body localization in Lig-depleted oocytes by approximately 1.3-1.5 fold (Fig. 7C-E). In contrast, *poe* and *ctrip* mRNA did not show a significant increase in P-body localization after Lig depletion (Fig. 7F-G), consistent with a lack of an effect of Lig knockdown on Poe protein levels. Bulk poly(A) RNA showed no comparable redistribution (Fig. 7H), indicating that Lig does not globally reorganize maternal mRNAs relative to P-bodies. Thus, Lig acts on a subset of FMRP targets by promoting protein synthesis and reducing P-body association of their transcripts, similar to FMRP’s observed activity.

## DISCUSSION

Our findings support a model in which FMRP safeguards long, dosage-sensitive mRNAs by controlling their access to competing cytoplasmic RNP states. Long FMRP targets appear particularly vulnerable to excessive Me31B/DDX6-dependent repression, potentially because their extended coding sequences provide large platforms for cooperative assembly of repressive RNP complexes. FMRP counteracts this vulnerability by maintaining target mRNAs in translation-competent states and limiting their shift into Me31B-marked P-body-associated repressive states. Thus, FMRP does not simply repress or activate translation in isolation; it controls partitioning of long target mRNAs between competing cytoplasmic RNP states.

The discrepancy between our findings and others supporting a repressive function for FMRP may reflect differences between the biochemical activities of isolated or inactive RNA-binding proteins and their functions within physiological RNP complexes. Indeed, other RGG-domain-containing proteins inhibit translation through interactions with initiation factors that occur *in vitro*^76^, yet fail to repress reporter expression when artificially tethered to mRNAs *in vivo*.^77^ Moreover, the RGG domain containing protein Npl3 was subsequently shown to promote translation initiation *in vivo*, directly contrasting with its inhibitory activity *in vitro*.^78^ These observations raise the possibility that inhibitory interactions mediated by arginine-rich or other low complexity domains, including electrostatic interactions that may be favored under cell-free assay conditions, do not necessarily reflect their dominant physiological activities. Rather, the translational effects of these proteins may depend on their interactions with other RNP components and the cellular environment that may not be replicated *in vitro* or in cell culture (Fig. 3).

The identification of Rin/G3BP, Lig/UBAP2L, and Capr/Caprin1, conserved core factors involved in stress granule assembly, as FMRP-associated cofactors connects this mechanism to a broader literature on cytoplasmic RNP organization and translational control. Although stress granules and P-bodies have traditionally been viewed as destinations for translationally inactive mRNAs,^79^ these assemblies are compositionally and functionally distinct. Stress granules contain mRNAs together with translation initiation factors and 40S ribosomal subunits, whereas P-bodies are enriched for decapping and repression machinery.^41,42,80,81^ Single-molecule imaging has further demonstrated that mRNAs within stress granules can undergo complete translation cycles and that translating transcripts can enter, exit, or remain within these assemblies.^82^ Consistent with this more active view, *Drosophila* Rin/G3BP and human UBAP2L promote the stability and translation of their bound mRNAs, while mammalian G3BP1/2-containing stress granules can weakly prioritize translation of stress-resistant transcripts during the integrated stress response.^43,73,83^ The potential translation-supportive functions of stress granule-associated machinery may be obscured by its study in cultured cells under conditions of acute stress, which dramatically reduces translation initiation. Developmentally specialized cells, including oocytes and neurons, must maintain extensive stores of translationally repressed mRNAs while preserving sufficient translation capacity for cellular functions. Indeed, studies of *Drosophila* germ granules have established relief of repression as an important mechanism of localized mRNA activation.^84,85^ In this context, our findings suggest that FMRP cooperates with core stress granule-associated factors to maintain sufficient target expression by preventing excessive recruitment into Me31B/DDX6-dependent repressive RNP states. More broadly, these observations raise the possibility that mRNA partitioning between distinct RNP environments may actively determine translational fate, with stress granule-associated machinery favoring translational competence and P-body-associated machinery favoring repression. Such organization may be particularly important for maintaining stored mRNA programs that underlie developmental competence and cellular plasticity.

This framework also places RNA length at the center of FMRP biology and may help explain why FMRP is especially important in neurons. The evolution of complex nervous systems was accompanied by the expansion of large, isoform-rich neuronal and synaptic genes, which increased the regulatory potential and isoform diversity of neuronal gene expression.^86,87^ Our findings suggest that this transcript architecture also creates a post-transcriptional challenge: long mRNAs require mechanisms that preserve their access to productive translation despite their increased exposure to abundant, low-specificity repressors such as Me31B/DDX6. FMRP may represent one such safeguard. In this view, the neuronal requirement for FMRP may arise not only from neuron-specific signaling or localization programs, but from the unusually long and dosage-sensitive mRNA landscape that neurons must express.

These findings reframe the consequences of FMRP loss as a failure of cytoplasmic RNP-state control. FMRP loss has often been interpreted through downstream neuronal phenotypes, including altered synaptic signaling, plasticity, and circuit development.^15,88^ Our data suggest that many phenotypes arise from an upstream failure to maintain long, dosage-sensitive neurodevelopmental mRNAs in translation-competent states. This model helps explain why loss of FMRP can reduce expression of large neurodevelopmental regulators despite the historical classification of FMRP as a repressor.^4,5,7,8,23^ Importantly, several FMRP targets encode proteins that themselves restrain gene-expression, signaling, proteostatic, or metabolic programs, including HUWE1, UBR4, and NF1/neurofibromin.^8,89-91^ Thus, failure to express FMRP targets may not simply deplete individual effector proteins; it may remove high-level regulators that normally restrain signaling, proteostasis, metabolism, and gene-expression programs. In this view, fragile X syndrome may reflect, in part, an upstream failure of RNP-state control that selectively compromises long regulatory mRNAs and propagates through the developmental networks those mRNAs control.

## METHODS

### Fly stocks and husbandry

Fly stocks were maintained at 25 °C. Flies were fed a standard cornmeal–yeast–dextrose medium. Per litre of food, the medium contained 70.9 g D-glucose (dextrose), 48.5 g cornmeal, 30.3 g active dry yeast, 20.5 g sucrose, 4.55 g agar, 0.5 g CaCl□·2H□O and 0.5 g MgSO□·7H□O, cooked in ∼0.47 L water and brought to final volume. After cooling, a propionic/phosphoric acid preservative mix was added to a final concentration of 0.49% (v/v) propionic acid and 0.049% (v/v) phosphoric acid. Detailed experimental genotypes and sources of fly stocks are listed in Supplementary Table S2.

### Cloning, gene editing and transgenesis

*Fmr1* and *Me31B* were endogenously tagged with 3xFLAG at their C-termini using CRISPR-mediated homologous recombination by Rainbow Transgenic Flies, Inc. In brief, tags were inserted by replacing the endogenous stop codon with a GS linker sequence, followed by a 3xFLAG tag, a stop codon, a synthetic SV40 3□UTR, and a sequence encoding the visible mini-white eye marker. *Poe* was similarly endogenously tagged at its C-terminus with GFP rather than 3xFLAG. The UASp-*me31B*-E208A construct was generated in the UASz1.0 backbone. A DNA fragment encoding the P-promoter sequence, an IVS intron, the Syn21 translational enhancer, and the full-length *me31B-RA* coding sequence carrying an E208A substitution in the conserved DEAD-box motif, fused in frame to EGFP via a flexible Gly–Ser linker (GSGGGGSGGGGS) and followed by the cyclin A (*cycA*) 3□UTR, was synthesized (GenScript) and inserted into the AatII and HpaI sites of UASz1.0. The construct was injected into attP40 landing-site lines (Genome Prolab Inc.) for □C31 integrase–mediated integration, and transformants were identified by screening for the mini-*white* (w□) eye-color marker.

Constructs encoding the RA isoform of full-length wild-type *Fmr1* (*Fmr1*^WT^) and the RNA-binding-deficient KH1 and KH2-domain mutant (*Fmr1*^KH1-KH2^ ^mut^) were generated in the UASz1.0 backbone using the same design as the *me31B*-E208A construct. A DNA fragment encoding the P-promoter sequence, an IVS intron, the Syn21 translational enhancer, and the full-length *Fmr1*-RA coding sequence, fused in frame at its C terminus to a 3xFLAG tag via a flexible Gly–Ser linker (GSGGGGSGGGGS), was synthesized and inserted into the NheI and BamHI sites of UASz1.0. To render the transgene resistant to *Fmr1* RNAi, synonymous mutations were generated in the RNAi-targeted region (coding sequence nucleotides 303 to 324) (TGAGACGTCCTACACCGAAATC to CGAAACCTCTTATACAGAGATA), which preserves the encoded amino acid sequence while disrupting complementarity to the shRNA (BDSC #35200). The mutant construct was identical except that the FMRP coding sequence additionally carried the I244N and I307N substitutions in the KH1 and KH2 RNA-binding domains, respectively. Constructs were injected and screened the same as the *me31B*-E208A transgenic flies.

### Enrichment of mature stage 14 oocytes

To obtain ovaries enriched for stage 14 oocytes, newly eclosed virgin females of the indicated genotypes were placed in standard food vials supplemented with yeast paste, prepared by mixing live yeast with water to a peanut-butter–like consistency that did not trap the flies. After 48 h of feeding, the flies were transferred to molasses vials containing agar–sugar medium (44 g agar, 180 mL molasses, 37 mL of 5% Tegosept, and 1,112 mL water) and maintained for a further 48 h; these vials provided humidity in the absence of edible yeast, thereby enriching for mature-stage (stage 14) oocytes. Unless otherwise specified, oocytes for all other experiments were collected from virgin females maintained on standard food supplemented with yeast paste for 1–2 days.

### RNA single molecule fluorescence *in situ* hybridization (smFISH)

smFISH was performed using reagents and protocol recommended by Biosearch Technologies™. In summary, fly oocytes were dissected in Grace’s insect medium and fixed in 4% paraformaldehyde (PFA) prepared in Grace’s insect medium for 12 minutes at room temperature with continuous agitation on a nutator. Following fixation, samples were washed three times in PBS supplemented with 0.1% Triton X-100 and 0.2% BSA (PBTB). Mature oocytes were subsequently bisected with a blade to enhance probe penetration. Samples were incubated in Buffer A for 20 minutes at room temperature, after which the buffer was replaced with hybridization buffer containing 10% formamide and incubated at 37 °C for 30–120 min. Custom RNA FISH probes (Supplementary Table S3) were designed using the Stellaris Probe Designer, diluted in hybridization buffer (1:50), and applied to the samples, which were then incubated overnight at 37 °C. Following hybridization, probe solution was removed and samples were incubated in Buffer A at 37 °C for 30 minutes, followed by an additional wash in Buffer A for 15 minutes at room temperature. Nuclei were stained with DAPI (5 mg/mL stock, diluted 1:20,000 in Buffer A) for 15 minutes at room temperature with agitation. Samples were then incubated in Buffer B for 10 minutes at room temperature with gentle agitation. Finally, tissues were mounted in 50% glycerol in PBS (30 μL per sample) and prepared for imaging. Slides were imaged on either a Leica SP8 confocal microscope with a 60× objective or an Olympus IXplore SR inverted spinning-disc fluorescence microscope with a 60× oil objective.

### Drug treatment

Wild-type (Oregon-R) ovaries were dissected into Grace’s insect medium supplemented with 100 μg/mL cycloheximide, 200 μM puromycin, or no additive (control), and incubated for 15 min on an orbital shaker at room temperature. Ovaries were then processed for single-molecule fluorescence *in situ* hybridization (smFISH) or immunostaining as described.

### Quantification of smFISH images

For the smFISH screen, *Poe* mRNA particles were quantified using Fiji/ImageJ (version 2.1.0). Nuclei identified by DAPI staining were masked out so that only the cytoplasmic *Poe* smFISH signal was carried into the downstream analysis. The cytoplasmic *Poe* signal was then segmented into two particle classes based on size and circularity (large and small). Large particles were defined as objects with an area of at least 0.2 μm² and a circularity of at least 0.5; small particles were defined from the signal remaining after large particles had been removed, as objects with an area below 0.2 μm² and a circularity of at least 0.4. The integrated intensity of each class was measured, and the fraction of *Poe* signal in large particles (integrated intensity of the large particles divided by the total *Poe* signal) was calculated and used to compare control fly lines against the RNAi and mutant lines in the screen.

smFISH signal from mature oocytes to detect *Fmr1* target mRNAs (*Poe, Ctrip, Huwe1, Nf1*, and *ana3*) or polyA RNA was quantified using a custom macro on Fiji/ImageJ (version 2.1.0). For each image, the smFISH and P-body channels were duplicated, split, and analyzed separately. To generate an RNA mask, the smFISH channel was background-subtracted using a rolling-ball radius of 100 pixels, corrected by subtracting a constant intensity value of 50, and manually thresholded using probe-specific settings. The thresholded image was converted to a binary mask, and RNA-positive objects were retained using Analyze Particles with a minimum size cutoff of 0.04 μm² and a circularity range of 0–1. The resulting mask was scaled to 0/1, converted to 16-bit, and multiplied by the background-subtracted smFISH image to generate an intensity-preserving RNA mask.

P-bodies were segmented from the P-body channel after background subtraction using a rolling ball radius of 200 pixels. P-body-positive pixels were identified using a fixed threshold determined from the mean P-body signal intensity and converted to a binary mask. This mask was scaled to 0/1 to define the P-body compartment, and an inverted mask was generated to define the non-P-body compartment. To quantify RNA signal associated with P-bodies, the intensity-preserving RNA mask was multiplied by either the P-body mask or the inverted P-body mask. Integrated density was measured in each resulting compartment to determine the amount of RNA signal inside and outside P-bodies.

### Immunostaining

Fly oocytes were dissected and fixed as described for the smFISH protocol above. Following fixation, samples were washed three times in PBTB for 10 minutes each with continuous agitation. Samples were then incubated with the primary antibodies rabbit anti-CAPR (1:500; laboratory of John C. Sisson, UT Austin), rabbit anti-POE (1:100; custom-generated by Proteintech), or mouse anti-FMR1 (1:1,000; clone 6A15, Abcam), diluted in PBST overnight at 4 °C. After primary antibody incubation, oocytes were washed three times in PBTB for 10 minutes per wash. Samples were then incubated overnight at 4 °C with Alexa Fluor-conjugated cross-adsorbed secondary antibodies diluted 1:1,000 in PBTB, selected according to the host species of the primary antibody and the fluorophore combination required for each experiment: goat anti-rabbit IgG (H+L) Alexa Fluor 568 (Invitrogen, A-11011), goat anti-rabbit IgG (H+L) Alexa Fluor 633 (Invitrogen, A-21070), goat anti-mouse IgG (H+L) Alexa Fluor 488 (Invitrogen, A-11001), and goat anti-mouse IgG (H+L) Alexa Fluor 568 (Invitrogen, A-11004). Following incubation, samples were washed three times in PBTB, with DAPI included in the final wash. After washing, 50% glycerol in PBS was added to tissues and mounted for imaging.

### Co-immunoprecipitation mass spectrometry (Co-IP/MS)

Fifty pairs of dissected fly ovaries were lysed in lysis buffer containing 150 mM NaCl, 5 mM MgCl□, 50 mM Tris-HCl (pH 7.5), 0.5% Triton X-100, and cOmplete™ Protease Inhibitor Cocktail (Roche #04693116001). Tissues were homogenized for 30 seconds on ice using an electric homogenizer. Lysates were centrifuged at 16,000 × g for 10 minutes at 4 °C, and supernatants were collected and either processed immediately or stored at −70 °C until use. Protein concentration was determined by BCA assay. For immunoprecipitation, anti-FLAG M2 magnetic beads (Sigma-Aldrich, M8823) were equilibrated twice in lysis buffer. A total of 500– 1000 μg of total protein lysate was incubated with equilibrated beads for 1 hour at room temperature with rotation. Beads were subsequently separated using a magnetic rack, and the supernatant was retained for assessment of depletion efficiency. Beads were washed three times with lysis buffer, and bound proteins were eluted by heat denaturation at 70 °C for 10 minutes in 2× Laemmli sample buffer containing 100 mM DTT. Eluted proteins were separated by SDS-PAGE using 4–15% Mini-PROTEAN® TGX™ precast gels. For mass spectrometry, gels were visualized with Coomassie stain; sections displaying protein bands were excised and submitted to the Proteomics Core Facility (UBC) for sample processing and LC-MS/MS. Proteins were reduced, alkylated, and digested with trypsin using an in-gel digestion protocol^92^.

Peptides were separated on a nanoflow UHPLC system (NanoElute 2, Bruker) and analyzed on a timsTOF Pro 2 mass spectrometer (Bruker) operated in DDA-PASEF mode. Spectra were searched against the *Drosophila melanogaster* UniProt database in Byonic (Protein Metrics) with a 1% protein-level false discovery rate. A total of six Co-IP/MS were performed from three *Fmr1*-FLAG replicates and three negative-control (Oregon-R) replicates.

### Cross-system analysis of FMRP binding and translation

FMRP binding maps were paired with measurements of translation after FMRP loss in four systems: *Drosophila* ovary (*Fmr1* iiCLIP, this study; translation efficiency in *Fmr1* RNAi vs control ovaries^4^), mouse cortex (P13 HITS-CLIP^93^; RiboDiff TE in P24 *Fmr1* KO vs WT cortex^23^), SH-SY5Y cells (FMRP-binding index recomputed from raw RIP-seq reads, DRA005644; SILAC protein minus RNA-seq log fold change in *FMR1* KO vs WT cells^58^) and HCT116 cells (*FMR1* CLIP from hPSC-derived forebrain neurons^6^; ribosome-footprint minus RNA log fold change in *FMR1* KO vs WT cells, base mean ≥ 10^94^). Cortex CLIP fragment counts were residualized on input RPKM to estimate binding per molecule. In each system, fold changes were divided by the median gene, transcripts were ranked by binding and split into ten equal-sized deciles, and the fractions decreased (< 0.8) or increased (> 1.2) were tested for trend with a two-sided Cochran–Armitage test, with Wilson 95% confidence intervals. The 200 most-bound transcripts were compared with all other transcripts by a two-sided Mann–Whitney U test. SFARI Gene class I and II genes (score 1–2, release 2026 Q2) among the 200 most-bound transcripts were identified by gene symbol in mammals and, in *Drosophila*, as reciprocal best DIOPT orthologs (score ≥ 5; FlyBase FB2026_03); decreased and increased genes were compared by a two-sided sign test. Analyses were performed in Python (v3.10) with pandas and SciPy.

### Normalization of spectral counts and interaction scoring

To correct for differences in total spectral output between affinity purifications, spectral counts were normalized prior to interaction scoring. For each immunoprecipitation, a size factor *s* was defined as the total spectral counts summed across all preys in that immunoprecipitation divided by the mean of these totals across all six immunoprecipitations. Every spectral count was then divided by the size factor of its immunoprecipitation and rounded to the nearest integer, equivalent to multiplying each count by (mean total)/(immunoprecipitation total). This scaled all six immunoprecipitations to a common total of approximately 7,220 spectra (raw totals ranged from 2,570 to 14,007; size factors ranged from 0.36 to 1.94). High-confidence interactors were identified with SAINTexpress in spectral-count mode (SAINTexpress-spc v3.6.1)^95^, using the three *Fmr1* immunoprecipitations as test runs and the three control immunoprecipitations as negative controls, with default parameters. Interactions with a Bayesian false discovery rate (BFDR) ≤ 0.05 were considered significant.

### Improved individual nucleotide resolution crosslinking immunoprecipitation sequencing (iiCLIP-seq) library preparation

The iiCLIP-seq protocol was adapted from Lee et al.^57^ with modifications optimized for *Drosophila* oocytes. Mature oocyte-enriched ovaries were dissected in Grace’s insect medium and flash-frozen in liquid nitrogen. Frozen tissue was pulverized to a fine powder using an electric mortar and pestle while submerged in liquid nitrogen and subjected to UV crosslinking at 254 nm using a GS Gene Linker® UV chamber (four exposures of 250 mJ/cm²). Crosslinking conditions were empirically optimized for each protein, and the minimal exposure yielding > 70% of maximal signal was used, in accordance with published recommendations.^57^

Tissue was lysed under denaturing conditions in buffer containing 50 mM Tris-HCl (pH 7.4), 100 mM NaCl, 1% Igepal CA-630, 2% SDS, 0.5% sodium deoxycholate, and protease inhibitors. Lysates were centrifuged at 16,000 × g for 10 minutes, and supernatants were incubated at 70 °C for 10 minutes with agitation (1,100 rpm). SDS concentration was subsequently reduced to 0.1%. Protein concentration was determined by BCA assay, and 1 mg of total protein was used for immunoprecipitation with ChromoTek DYKDDDDK Fab-Trap® agarose beads. Samples were incubated with beads for 1 hour at 4 °C with continuous rotation.

Following immunoprecipitation, beads were washed overnight in high-salt wash buffer (50 mM Tris-HCl, (pH 7.4), 1 M NaCl, 1 mM EDTA, 1% Igepal CA-630, 0.1% SDS, and 0.5% sodium deoxycholate). All subsequent steps, including library preparation and sequencing, were performed as described in Lee et al.^57^ without modification. Two biological replicate libraries were sequenced for each protein (FMRP-FLAG and Me31B-FLAG).

### iiCLIP-seq data analysis

Two biological replicates per protein were analyzed separately. Reads were demultiplexed, UMIs were extracted, and adapters were trimmed with cutadapt (v5.2), keeping inserts ≥ 18 nt. Reads were then aligned uniquely to the *D. melanogaster* BDGP6.54 genome (Ensembl release 116) with STAR (v2.7.10b) and deduplicated by UMI with UMI-tools (v1.1.6). Crosslink sites were defined as the nucleotide upstream of each read’s 5□ end and restricted to protein-coding genes. They were assigned to the 5□UTR, CDS or 3□UTR using iCount-Mini (v4.0.1) segmentation. Binding was quantified as exonic crosslink counts per million divided by the mean RNA-seq transcripts per million (TPM) from oocyte RNA-seq. This was defined as crosslink sum. Dividing the crosslink sum by mRNA length in kb gave crosslink density. Values were averaged across replicates for the 6,152 mRNAs with TPM ≥ 5 and ≥ 1 crosslinks in every library. High-binding targets were mRNAs above the knee of the ranked crosslink sum curve. For length and expression analyses, mRNAs were grouped into equal-sized bins, and bin medians were normalized to the first bin, with 95% bootstrap confidence intervals (10,000 resamplings). For comparison with mouse^93^ and human^6^ FMRP CLIP, binding was converted to percentiles within each dataset and related to CDS length by Spearman correlation. SFARI class I–II orthologs were defined as reciprocal best DIOPT matches (score ≥ 5); their enrichment across binding deciles was tested with the Cochran–Armitage trend test. Translation efficiency changes in *Fmr1* RNAi oocytes were obtained from Flanagan et al.^8^. The enrichment of targets with reduced TE (adjusted *p* < 0.01) was tested with a two-sided Fisher’s exact test, and overlap between FMRP and Me31B targets with a hypergeometric test.

### Western blotting

Mature-stage oocytes were enriched as described above (see Enrichment of mature-stage oocytes). Ovaries were dissected in room-temperature Grace’s Insect Medium and lysed in lysis buffer (0.5% Triton X-100, 150 mM NaCl, 5 mM MgCl□, 50 mM Tris-HCl pH 7.5, 20 U/mL SUPERase·In [Ambion], and 40 mM NEM [Sigma-Aldrich]). Protein concentration was determined by BCA assay, and samples were loaded onto 4–15% Mini-PROTEAN® TGX Stain-Free™ gels (Bio-Rad, #4568085) and electrophoresed at 130 V for 1 h. Gels were activated and imaged for total protein on a Bio-Rad ChemiDoc imaging system. Proteins were transferred onto PVDF membranes in Towbin buffer at 350 mA for 3–4 h (4 h for POE; 3 h for VPS13 and ANA3). Membranes were blocked in Odyssey Blocking Buffer (LI-COR) for 30–60 min at room temperature, then incubated overnight at 4 °C with gentle agitation in Odyssey Blocking Buffer containing 0.1% Tween-20 with the primary antibodies rabbit anti-POE (1:100; custom-generated by Proteintech), rabbit anti-VPS13 (1:1,000; laboratory of Ody C.M. Sibon, UMCG), or rabbit anti-ANA3 (1:1,000; laboratory of David Glover, Caltech). Membranes were washed three times in PBST (PBS with 0.1% Tween-20) and incubated with IRDye secondary antibodies (1:15,000; LI-COR) for 1 h at room temperature. Membranes were then washed three times in PBST and twice in PBS for 5 min per wash, and imaged using a ChemiDoc Imaging System (Bio-Rad). The same procedure was used to detect FLAG-tagged immunoprecipitation eluates, using the primary antibodies mouse anti-FLAG (1:5,000; F3165 [Sigma-Aldrich]), mouse anti-ME31B (1:5,000; laboratory of Akira Nakamura, Kumamoto University), rabbit anti-RIN (1:8,000; laboratory of Éric Lécuyer, IRCM), and rabbit anti-CAPR (1:3,000; laboratory of John C. Sisson, UT Austin).

### FMRP-FLAG co-immunoprecipitation

For co-immunoprecipitation followed by Western blotting, ovaries were dissected from 30 well-fed females, and lysates were prepared and immunoprecipitated with FLAG magnetic agarose beads as described above (see Co-immunoprecipitation mass spectrometry), with the following modifications. A total of 2 mg of whole-ovary protein lysate was used per immunoprecipitation. After the final wash, bound proteins were eluted by competitive elution in 500 μg/mL 3x DYKDDDDK peptide (Pierce, Thermo Scientific, #A36805) for 10 min at room temperature with gentle agitation, and the eluate was then denatured by heating in 2× Laemmli sample buffer containing 100 mM DTT. The eluate was divided equally and loaded across two gels, with each gel receiving half of the eluate (corresponding to 1 mg of input lysate), alongside an input sample corresponding to 1.5% of the lysate used for the immunoprecipitation. One blot was probed for FLAG and CAPR, and the other for ME31B and RIN.

### Ribosome profiling and mRNA-seq library preparation

Ribosome footprinting was performed as described^4,96^ with several modifications. Forty pairs of ovaries enriched for stage 14 mature stage oocytes were isolated through dissection in Grace’s insect medium (Gibco^TM^, #11605094) and flash frozen in liquid nitrogen. Ovaries were lysed in 200 μL of ribosome profiling lysis buffer (0.5% Triton X-100, 150 mM NaCl, 5 mM MgCl□, 50 mM Tris-HCl pH 7.5, 1 mM DTT, 20 μg/mL emetine [Sigma-Aldrich], 20 U/mL SUPERase·In [Ambion], and 50 μM GMP-PNP [Sigma-Aldrich]) by grinding for 15 s with a plastic micropestle. Total RNA was quantified using the Qubit RNA Broad Range Assay Kit, and the *Drosophila melanogaster* lysate (130–140 μg of total RNA) was spiked with a whole-ovary lysate from *Drosophila pseudoobscura* corresponding to 2% of total RNA (w/w) prior to nuclease digestion, such that *D. pseudoobscura* ribosomes were footprinted in parallel as an external normalization standard. Ribosome footprints were generated by incubating the combined lysate with micrococcal nuclease S7 (Roche) at 2 U per μg of total RNA for 40 min at 25 °C, after which the reaction was quenched by adding EGTA to a final concentration of 6.25 mM. Ribosomes were sedimented through a 34% sucrose cushion by centrifugation at 80,000 rpm for 1 h 40 min in a TLA-110 rotor, and the resulting pellet was resuspended in 10 mM Tris-HCl (pH 7.0). RNA was extracted from monosomes by TRIzol (Invitrogen) and size-selected on a 10% TBE–urea gel. Size-selected footprints were dephosphorylated, and library preparation and ribosomal RNA depletion were performed as described^96^, using the *Drosophila* rRNA depletion oligonucleotides reported in Dunn et al.^97^.

For mRNA-seq, 10 pairs of stage-14–enriched ovaries were dissected and lysed directly in 1 mL of TRIzol™ reagent (Invitrogen) and homogenized for 15 s with a plastic micropestle. After incubation at room temperature for 5 min, lysates were centrifuged at 12,000 × g for 10 min at 4 °C. Supernatants were transferred to a fresh tube, 200 μL of chloroform was added, and samples were incubated for 3 min at room temperature before centrifugation at 10,000 × g for 15 min at 4°C. The upper aqueous phase was transferred to a fresh tube, 0.6 mL of isopropanol was added, and samples were incubated for 10 min at room temperature. RNA was precipitated by centrifugation at 20,000 × g for 20 min at 4 °C; the pellet was washed with 1 mL of ice-cold 80% ethanol and resuspended in water. Total RNA was quantified using the Qubit RNA Broad Range Assay Kit. Spike-in RNA was prepared in parallel by direct TRIzol extraction of whole *D. pseudoobscura* ovaries. Immediately before poly(A) selection, 250 ng of total RNA containing a 5 ng (2%) *D. pseudoobscura* RNA spike-in was used for mRNA enrichment with NEBNext Oligo d(T)□□ beads (NEB, #S1419S), and libraries were prepared using the NEBNext UltraExpress® RNA Library Prep Kit (NEB, #E3330S) according to the manufacturer’s instructions. For the experiments shown in Figure S5, mRNA-seq was performed in three biological replicates each for *Rin* null, *lig* knockdown, and *Capr* null ovaries. mRNA-seq data for *Fmr1*-depleted ovaries were obtained from Greenblatt and Spradling^4^. For the experiments shown in Figure 4 and Figure S3, ribosome profiling and mRNA-seq were each performed in three biological replicates for each of the four conditions, yielding 24 libraries in total.

### Ribosome profiling and mRNA-seq data analysis

Analyses of ribosome profiling and mRNA sequencing data were conducted as in Flanagan et al.^8^, with the following modifications. Both ribosome profiling and mRNA-seq libraries were sequenced as 150-bp paired-end reads and read 1 was used for all analyses. Reads were processed with the FASTX-Toolkit. For ribosome profiling, adapters were removed with fastx_clipper (-Q33 -a AAAAAAAA -l 25 -c -n), retaining only adapter-containing reads of ≥25 nt, after which the first three nucleotides of each read were trimmed with fastx_trimmer (-f 4); noncoding-RNA–derived reads, including rRNA, were then depleted *in silico* by alignment to the *D. melanogaster* noncoding-RNA reference (Ensembl BDGP6.32 ncRNA) using Bowtie2 (-L 23), and the unaligned reads were carried forward. For mRNA-seq, no adapter trimming was required because the insert sizes exceeded the read length, and no rRNA-depletion step was applied because the libraries were poly(A)-enriched.

For quantification of bulk mRNA and ribosome-footprint levels, the resulting reads from both assays were aligned with STAR (v2.7.9a) to a reference comprising the combined coding sequences of *D. melanogaster* and *D. pseudoobscura* (NCBI RefSeq assembly GCF_009870125.1, UCI_Dpse_MV25; cds_from_genomic), retaining only uniquely mapping reads (-- outFilterMultimapNmax 1). Reads mapping to *D. melanogaster* and to *D. pseudoobscura* were tabulated (Table S4), with the *D. pseudoobscura* counts serving as the external spike-in normalization standard. Uniquely mapping *D. melanogaster* reads were then re-aligned to the *D. melanogaster* genome (Release 6.32) with STAR (v2.7.9a), and read counts were assigned with featureCounts (Subread, v2.0.1) using the FlyBase dmel-all-r6.49.gtf annotation over coding sequences.

### Differential translation-efficiency (□TE) analysis

Per-gene featureCounts outputs were assembled into a single count matrix for downstream analysis. Global translation efficiency (Figure 4B) was computed independently of the DESeq2 model. Ribosome-profiling and mRNA-seq libraries were prepared from independent ovary collections. For each library, the ratio of *D. melanogaster* to *D. pseudoobscura* reads was computed and corrected for preparation batch by multiplying by the ratio of the overall geometric mean to the batch geometric mean, calculated separately within each assay. Batches corresponded to: for ribosome profiling, replicate 1 and replicates 2 to 3; for mRNA-seq, replicates 1 to 2 and replicate 3. Because the assays are unpaired, translation efficiency was calculated at the genotype level as the geometric mean of the corrected ribosome-footprint ratios divided by the geometric mean of the corrected mRNA ratios, expressed relative to control oocytes. Significance was assessed by a two-sided permutation test in which genotype labels were permuted independently within each assay; all 20 × 20 = 400 possible assignments were enumerated, giving a minimum attainable *p*-value of 0.005.

For enrichment analysis (Figure 4C and Figure 4C□), the per-gene □TE results were classified as significantly changed at adjusted *p* < 0.01 and split by direction of change. High-confidence binding targets were defined by iiCLIP score, using thresholds of >25 for FMRP and >15 for Me31B. For each target set (FMRP targets, n = 216; Me31B targets, n = 354), the proportion of significantly up- or downregulated genes was compared with the non-target background using a two-sided Fisher’s exact test, performed separately for each contrast and each direction.

Per-gene changes in translation efficiency (Figure 4D and 4E) were analyzed in R (v4.2.2) with DESeq2 (v1.38.3)^98^. Changes in translation efficiency were modeled with the interaction design ∼ batch + assay + condition + assay:condition, in which the assay:condition interaction term tests condition-dependent changes in the ribosome-footprint-to-mRNA ratio relative to control oocytes. Because libraries were prepared in separate sessions, a three-level batch factor was included in the model, grouping samples as normal (mRNA-seq replicates 1 and 2 and ribosome-profiling replicate 1), the mRNA-seq replicate-3 outlier, and the ribosome-profiling replicate 2 and 3 outlier. Size factors were set from the *D. pseudoobscura* spike-in: per-sample spike-in read counts (Table S4) were divided by their geometric mean across all libraries and supplied to DESeq2 as custom size factors, so that normalization reflected spike-in abundance rather than total library depth. Low-count genes were removed by DESeq2 independent filtering, which is computed per contrast; consequently, the number of testable genes differs slightly among comparisons (6,501 to 6,532). The model was fit using DESeq2 default settings. Comparisons against control oocytes were read directly from the corresponding assay:condition interaction coefficient, whereas comparisons between two non-reference conditions were evaluated as Wald tests on the difference between the relevant interaction coefficients. *p*-values were adjusted for multiple testing by the Benjamini-Hochberg method, and genes with adjusted *p* < 0.01 were considered significant.

## Supporting information

Table S1_SFARI Gene enrichment within FMRP and Me31B targets

Table_S2_Genotypes of the exeprimental files

Table_S3_poe_smFISH_probes

Table_S4_spikein_normalization

Table_S5_ribosome_profiling_oligos

Table_S6_Key_resources_table

## ACKNOWLEDGEMENTS

We thank Dr. Flora C.Y. Lee for her guidance with the iiCLIP-seq protocol and Crystal Shan for help with the data analysis. We are grateful to Dr. Akira Nakamura (Kumamoto University) for the ME31B antibody, Dr. David Glover (California Institute of Technology) for the ANA3 antibody, Dr. Ody C.M. Sibon University Medical Center Groningen (UMCG), for the VPS13 antibody, Dr. John C. Sisson and Dr. Ophelia Papoulas (University of Texas at Austin) for the CAPR antibody, and Dr. Éric Lécuyer (Institut de recherches cliniques de Montréal) for the RIN antibody. We also thank Dr. Ming Gao (Indiana University Northwest) for the *me31B*-GFP fly line. This work was supported by the Canadian Institutes of Health Research (CIHR), the Simons Foundation Autism Research Initiative (SFARI), Michael Smith Health Research BC, and the Lillian Lincoln Foundation.

## DECLARATION OF AI-ASSISTED TECHNOLOGIES

Claude (Anthropic) and ChatGPT (OpenAI) were used to edit author-written text for grammar, clarity and conciseness, and to assist in writing and debugging analysis scripts for ribosome profiling analysis and iiCLIP-seq data analysis. All text and code were inspected and validated by the authors. No large language model was used to generate, analyze or interpret experimental data, or to produce any figure.

## SUPPLEMENTAL MATERIAL

### TABLE LEGENDS

**Table S1. SFARI Gene enrichment within FMRP and Me31B targets**

**Table S2. Genotypes and sources of the *Drosophila* lines used in this study.**

**Table S3. Stellaris smFISH probe set for *poe*.**

**Table S4. *Drosophila pseudoobscura* spike-in read counts used for library normalization.**

**Table S5. *Drosophila melanogaster* rRNA depletion oligonucleotides used in Ribosome profiling**

**Table S6. Key resources table**

### FIGURES

**Figure S1.**
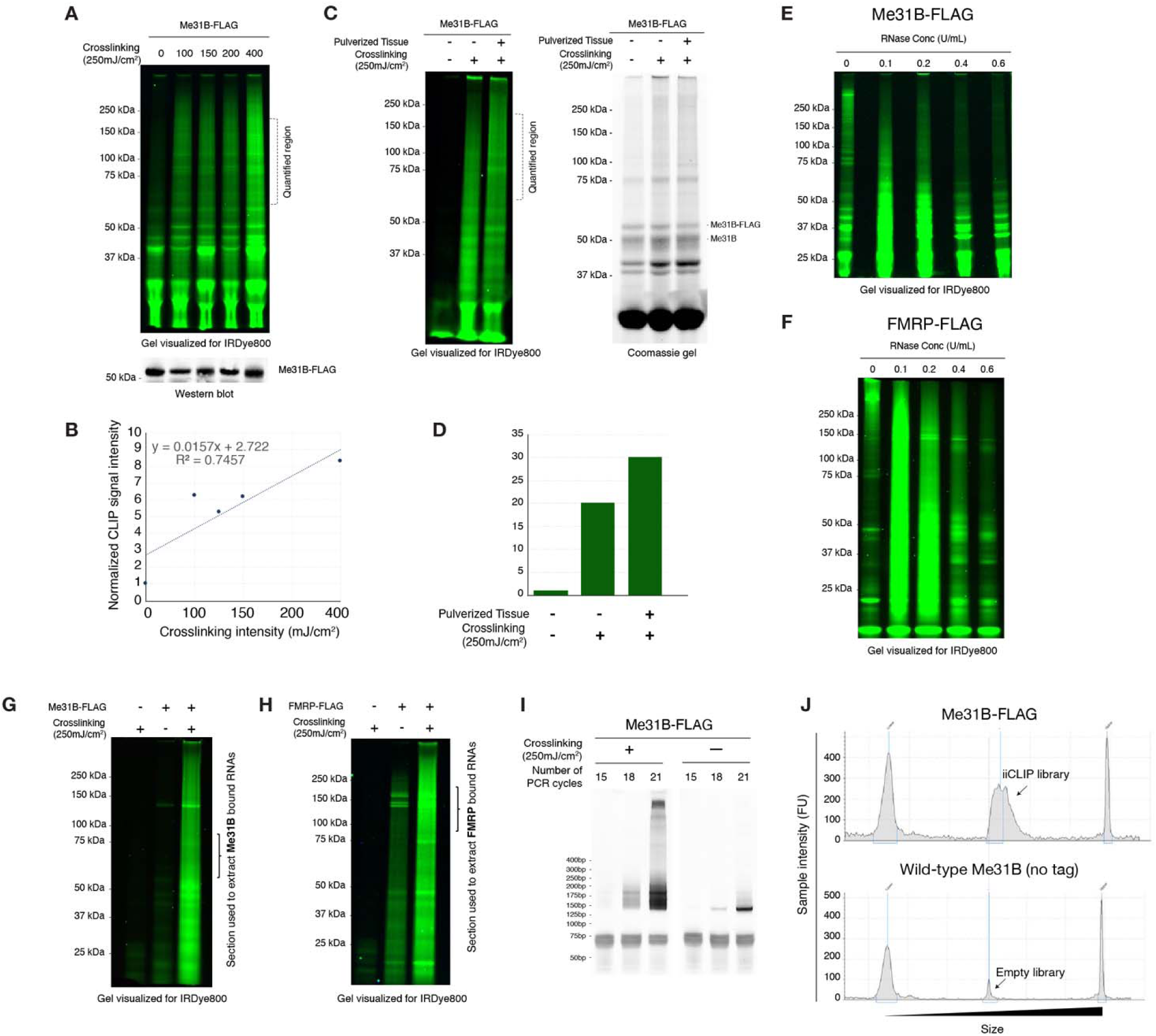
iiCLIP-seq optimization in *Drosophila* oocytes. **(A)** Optimization of UV crosslinking intensity showing increasing crosslinked RNA signal at higher intensities. Signal on gel is fluorescence from IRDye800 conjugated adapter ligated to RNAs that co-immunoprecipitated with Me31B-FLAG. Western blot of Me31B-FLAG showing protein levels following co-immunoprecipitation (bottom). **(B)** Quantification of **(A)**. **(C)** Comparison of crosslinking efficiency in flash frozen, pulverized oocyte tissue versus intact tissue suspended in Grace’s media (left) and Coomassie gel showing co-immunoprecipitated Me31B-FLAG indicating no effect on protein degradation in intact versus pulverized tissue (right). **(D)** Quantification of **(C)**. **(E-F)** Optimization of RNase concentration used prior to protein co-immunoprecipitation for Me31B-FLAG **(E)** and FMRP-FLAG **(F)**. **(G-H)** Representative gel images showing crosslinked RNA signal to Me31B-FLAG **(G)** and FMRP-FLAG **(H)** with minimal background when co-immunoprecipitated using FLAG Fab. These samples were used for final library preparation. **(I)** Optimization of PCR cycles used for library preparation showing amplified cDNA library signal for crosslinked Me31B-FLAG but not for negative control. **(J)** Quantification of final libraries using Agilent TapeStation showing broad peak for Me31B-FLAG but not for un-tagged control.

**Figure S2.**
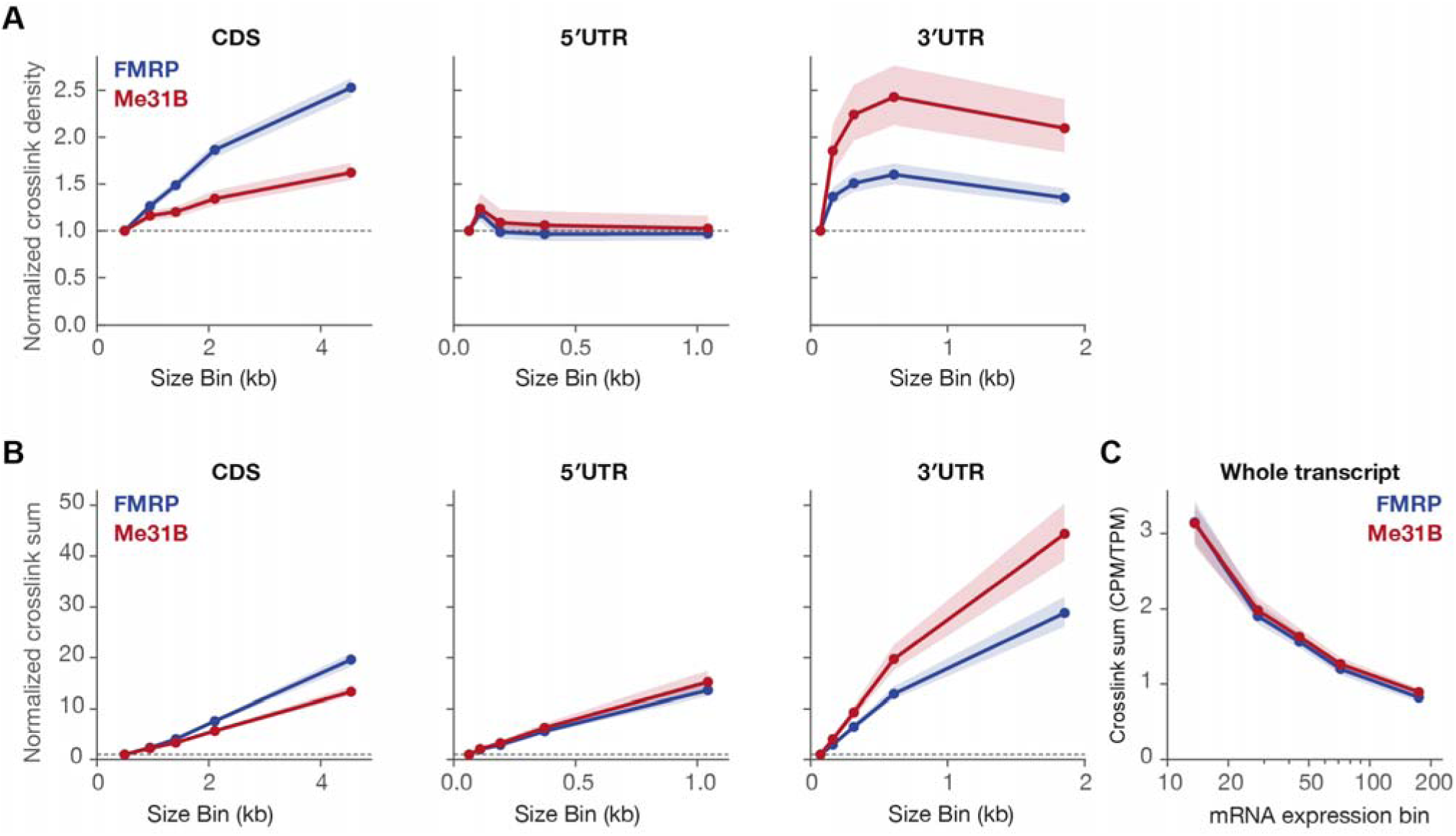
FMRP and Me31B bind mRNAs proportionally to CDS length and along similar motifs. **(A-B)** FMRP (blue) and Me31B (red) normalized crosslink density **(A)** and normalized crosslink sum **(B)** plotted along median mRNA length within increasing size bins showing that binding increases strongly with increasing length of CDS compared to UTRs. **(C)** FMRP (blue) and Me31B (red) peak sum plotted against RNA-seq bins showing that lowly expressed transcripts have higher binding by both proteins.

**Figure S3.**
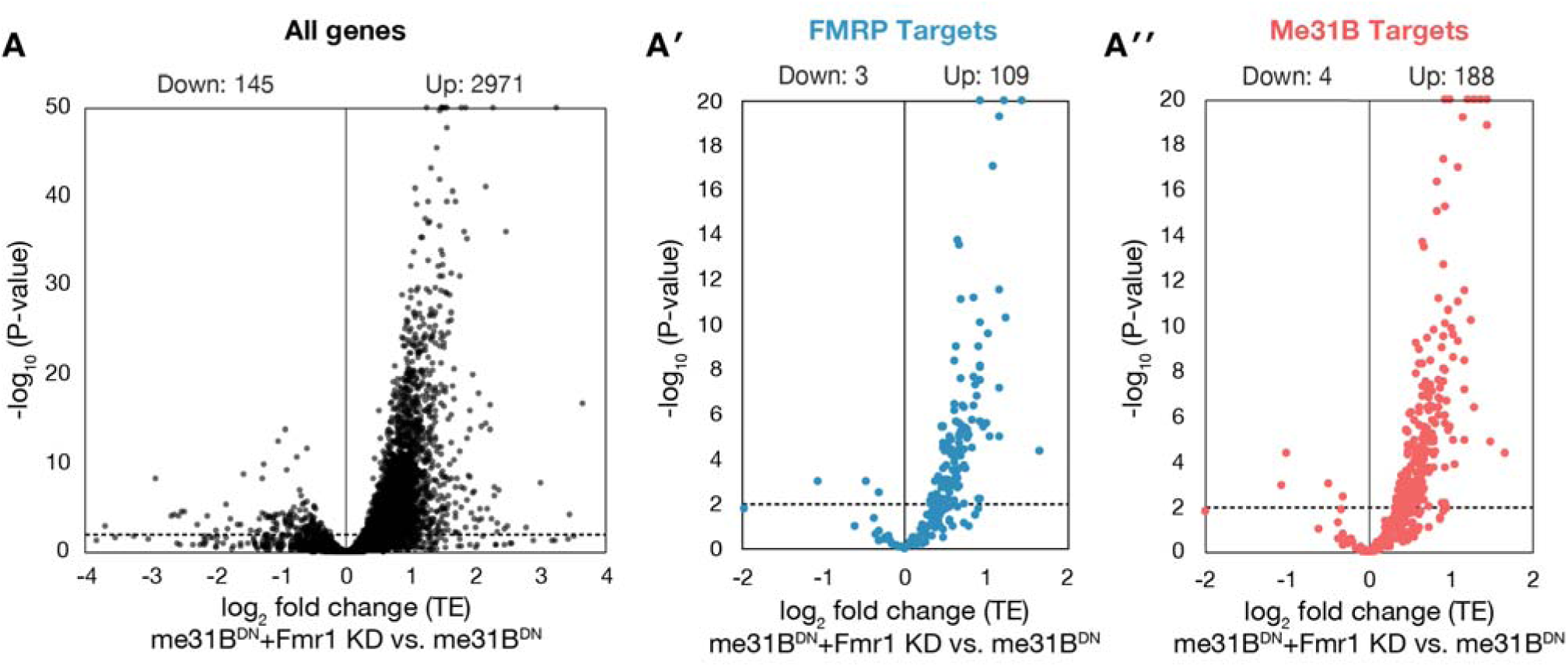
Translation efficiency changes upon *Fmr1* knockdown in the Me31B^DN^ background. Volcano plots of TE change in Me31B^DN^ plus *Fmr1* KD versus Me31B^DN^ for **(A)** all DESeq2-analyzed genes, **(A□)** FMRP targets, and **(A□)** Me31B targets, showing a pronounced positive shift in log_2_ TE fold change for both target sets. In all volcano plots the dashed line marks the significance threshold (adjusted *p* < 0.01); the x-axis is log_2_ TE fold change and the y-axis is −log_10_ adjusted *p*, and Up and Down gene counts are indicated above each plot.

**Figure S4.**
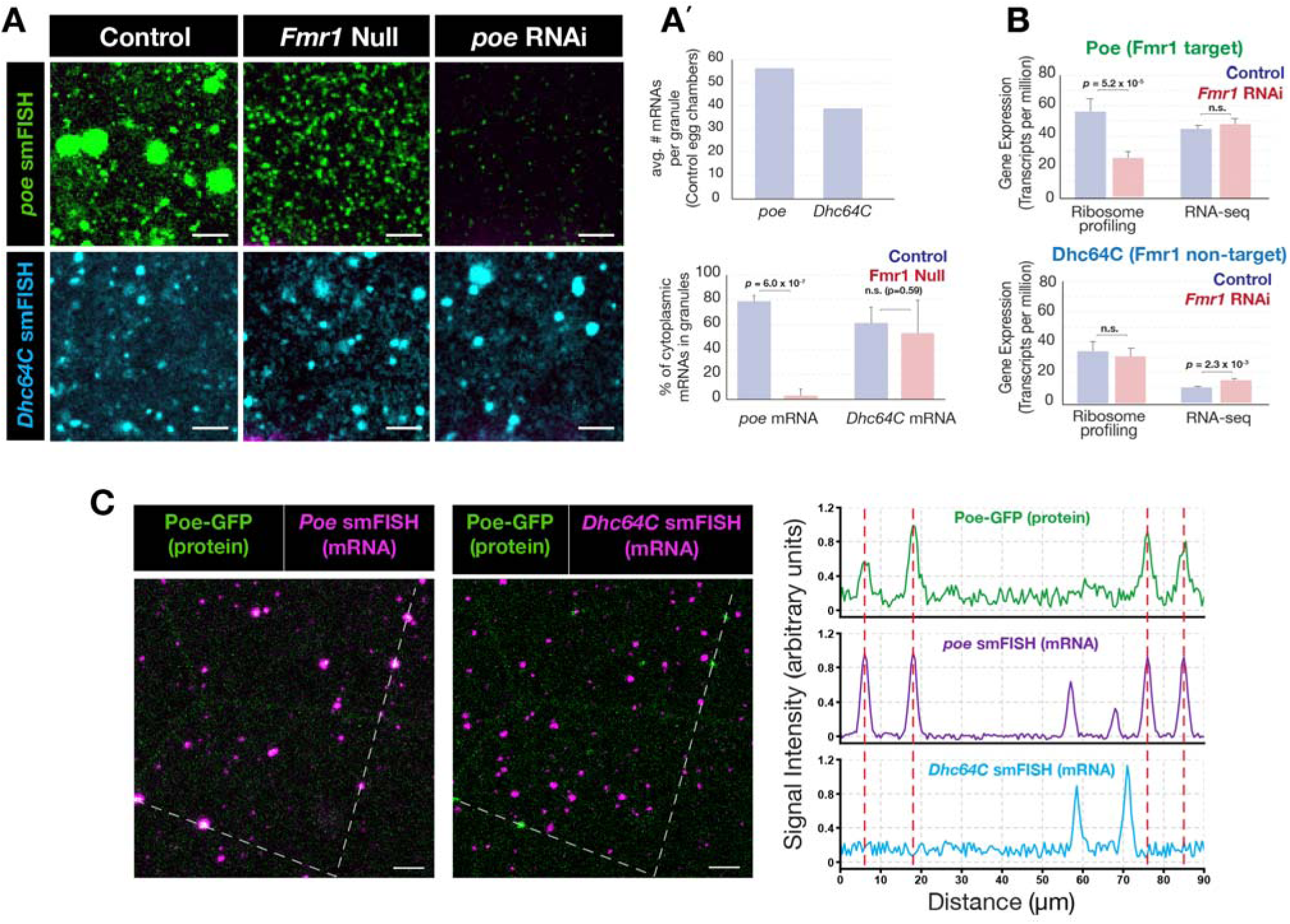
*poe* particle formation is target-specific and *poe* mRNA colocalizes with full-length Poe protein. **(A)** *poe* smFISH (top) and *Dhc64C* smFISH (bottom) in control, *Fmr1*-null (*Fmr1³/Fmr1*□^50^), and *poe* RNAi stage 10 nurse cells. **(A□)** Top, quantification showing an average of ∼55 mRNAs per *poe* granule; bottom, the fraction of cytoplasmic mRNA in particles is strongly reduced for *poe*, an FMRP target, in *Fmr1*-null follicles (*p* < 0.01) but unchanged for *Dhc64C*, an FMRP non-target (n.s., *p* = 0.59), indicating that loss of particle localization is specific to the *Fmr1* target *poe*. **(B)** Plots showing that the translation of Poe mRNA, but not that of another large mRNA, *Dhc64C*, is reduced in *Fmr1* RNAi oocytes. **(C)** Full-length Poe-GFP protein with *poe* smFISH (left) or *Dhc64C* smFISH (middle) in late wild-type follicles, with line scans at right showing that *poe* mRNA peaks coincide with Poe-GFP protein peaks whereas *Dhc64C* mRNA peaks do not, showing that mRNA-protein coincidence in these particles is specific to *poe*.

**Figure S5.**
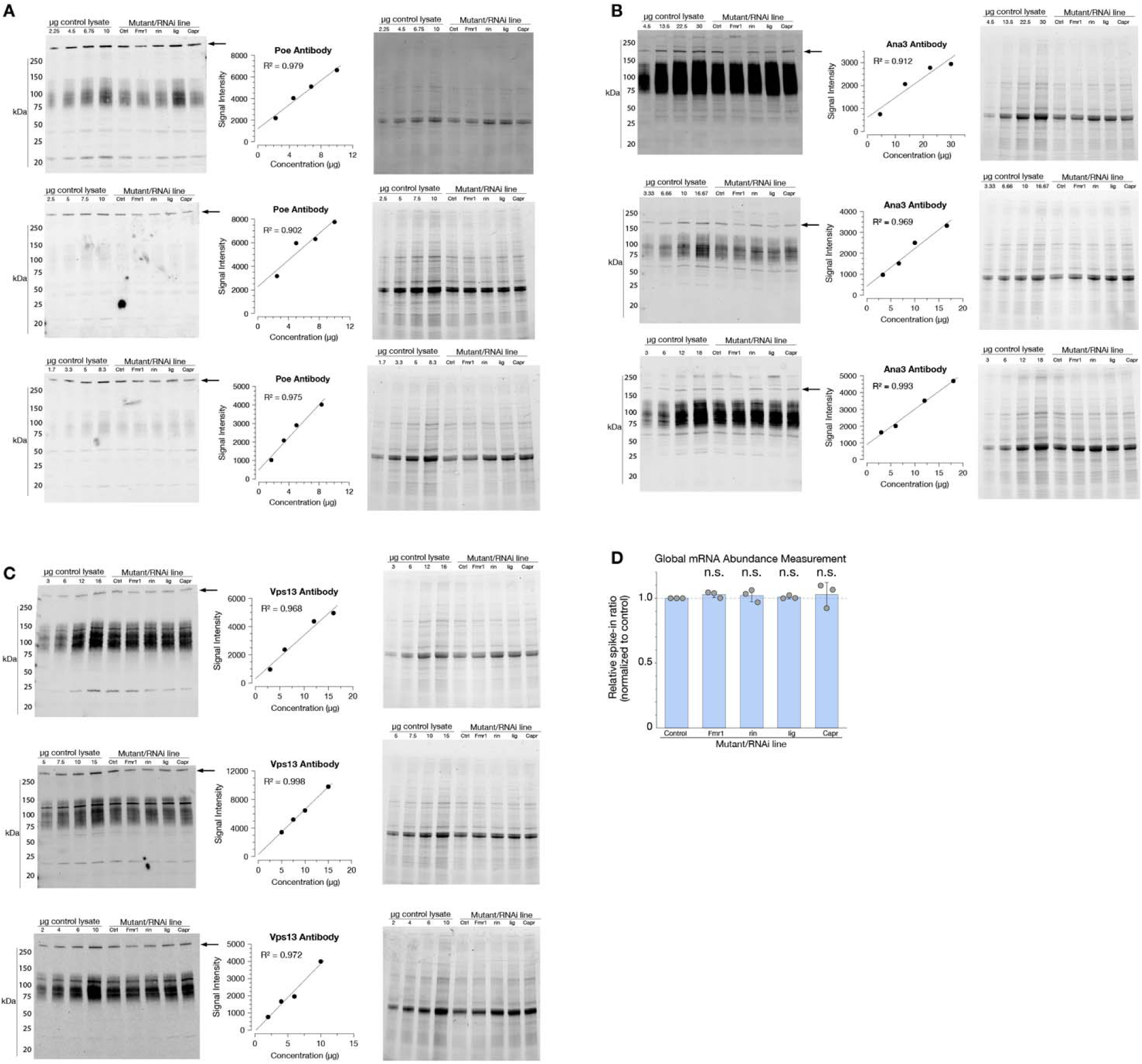
Quantitative Western blot measurement of FMRP target protein levels across FMRP-associated cofactor perturbations. **(A–C)** Full scans of Western blots (left) from oocyte extracts of the indicated control (Ctrl) and *Fmr1*, *rin*, *Capr*, and *lig* mutant or RNAi genotypes, probed for the *FMR1* targets **(A)** Poe, **(B)** Ana3, and **(C)** Vps13, each loaded with a control-lysate dilution series for quantification, together with the linearity of Western blot signal for each antibody (middle; *R*² ≥ 0.90) and the stain-free total-protein gel used for total protein normalization (right). **(D)** Chart showing that global mRNA abundance, measured as the spike-in-normalized RNA ratio relative to control, is unchanged across *Fmr1*, *rin*, *lig*, and *Capr* genotypes (n.s.), indicating that the reductions in Poe, Ana3, and Vps13 protein are not accounted for by loss of total target mRNA.

**Figure S6.**
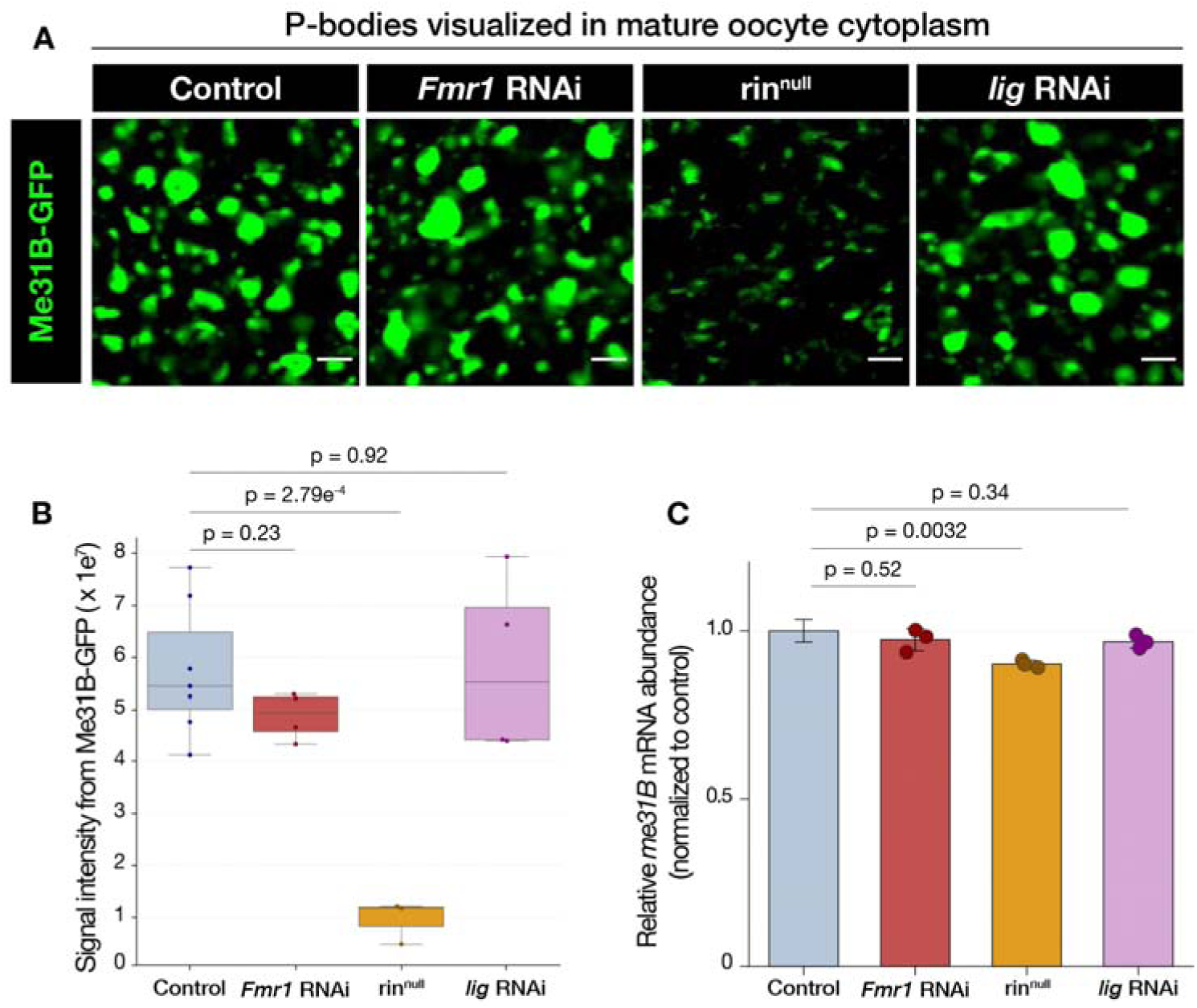
P-bodies remain unaffected by depletion of FMRP and Lig but not Rin. **(A)** Me31B-GFP in mature oocyte cytoplasm representing P-bodies in Control (*mCherry* RNAi), *Fmr1* RNAi, rin^null^, and *lig* RNAi. Scale bars: 3 μm. **(B)** Quantification of Me31B-GFP signal from **(A)**. **(C)** *me31B* mRNA abundance measured by RNA-seq shows that its mRNA levels decrease significantly in rin^null^ oocytes but not in control or *Fmr1* and *lig*-RNAi expressing oocytes.

## REFERENCES

1. King, I.F., Yandava, C.N., Mabb, A.M., Hsiao, J.S., Huang, H.S., Pearson, B.L., Calabrese, J.M., Starmer, J., Parker, J.S., Magnuson, T., et al. (2013). Topoisomerases facilitate transcription of long genes linked to autism. Nature 501, 58–62. 10.1038/nature12504.

2. Gabel, H.W., Kinde, B., Stroud, H., Gilbert, C.S., Harmin, D.A., Kastan, N.R., Hemberg, M., Ebert, D.H., and Greenberg, M.E. (2015). Disruption of DNA-methylation-dependent long gene repression in Rett syndrome. Nature 522, 89–93. 10.1038/nature14319.

3. Iossifov, I., O’Roak, B.J., Sanders, S.J., Ronemus, M., Krumm, N., Levy, D., Stessman, H.A., Witherspoon, K.T., Vives, L., Patterson, K.E., et al. (2014). The contribution of de novo coding mutations to autism spectrum disorder. Nature 515, 216–221. 10.1038/nature13908.

4. Greenblatt, E.J., and Spradling, A.C. (2018). Fragile X mental retardation 1 gene enhances the translation of large autism-related proteins. Science 361, 709–712. 10.1126/science.aas9963.

5. Sawicka, K., Hale, C.R., Park, C.Y., Fak, J.J., Gresack, J.E., Van Driesche, S.J., Kang, J.J., Darnell, J.C., and Darnell, R.B. (2019). FMRP has a cell-type-specific role in CA1 pyramidal neurons to regulate autism-related transcripts and circadian memory. eLife 8, e46919. 10.7554/eLife.46919.

6. Li, M., Shin, J., Risgaard, R.D., Parries, M.J., Wang, J., Chasman, D., Liu, S., Roy, S., Bhattacharyya, A., and Zhao, X. (2020). Identification of FMR1-regulated molecular networks in human neurodevelopment. Genome Res. 30, 361–374. 10.1101/gr.251405.119.

7. Seo, S.S., Louros, S.R., Anstey, N., Gonzalez-Lozano, M.A., Harper, C.B., Verity, N.C., Dando, O., Thomson, S.R., Darnell, J.C., Kind, P.C., et al. (2022). Excess ribosomal protein production unbalances translation in a model of Fragile X Syndrome. Nat. Commun. 13, 3236. 10.1038/s41467-022-30979-0.

8. Flanagan, K., Baradaran-Heravi, A., Yin, Q., Dao Duc, K., Spradling, A.C., and Greenblatt, E.J. (2022). FMRP-dependent production of large dosage-sensitive proteins is highly conserved. Genetics 221, iyac094. 10.1093/genetics/iyac094.

9. Sullivan, S.D., Welt, C., and Sherman, S. (2011). FMR1 and the continuum of primary ovarian insufficiency. Semin. Reprod. Med. 29, 299–307. 10.1055/s-0031-1280915.

10. Hagerman, R.J., Berry-Kravis, E., Hazlett, H.C., Bailey, D.B., Jr., Moine, H., Kooy, R.F., Tassone, F., Gantois, I., Sonenberg, N., Mandel, J.L., et al. (2017). Fragile X syndrome. Nat. Rev. Dis. Primers 3, 17065. 10.1038/nrdp.2017.65.

11. Darnell, J.C., Van Driesche, S.J., Zhang, C., Hung, K.Y.S., Mele, A., Fraser, C.E., Stone, E.F., Chen, C., Fak, J.J., Chi, S.W., et al. (2011). FMRP stalls ribosomal translocation on mRNAs linked to synaptic function and autism. Cell 146, 247–261. 10.1016/j.cell.2011.06.013.

12. Abrahams, B.S., Arking, D.E., Campbell, D.B., Mefford, H.C., Morrow, E.M., Weiss, L.A., Menashe, I., Wadkins, T., Banerjee-Basu, S., and Packer, A. (2013). SFARI Gene 2.0: a community-driven knowledgebase for the autism spectrum disorders (ASDs). Mol. Autism 4, 36. 10.1186/2040-2392-4-36.

13. Korb, E., Herre, M., Zucker-Scharff, I., Gresack, J., Allis, C.D., and Darnell, R.B. (2017). Excess translation of epigenetic regulators contributes to Fragile X syndrome and is alleviated by Brd4 inhibition. Cell 170, 1209–1223. 10.1016/j.cell.2017.07.033.

14. Ascano, M., Jr., Mukherjee, N., Bandaru, P., Miller, J.B., Nusbaum, J.D., Corcoran, D.L., Langlois, C., Munschauer, M., Dewell, S., Hafner, M., et al. (2012). FMRP targets distinct mRNA sequence elements to regulate protein expression. Nature 492, 382–386. 10.1038/nature11737.

15. Richter, J.D., and Zhao, X. (2021). The molecular biology of FMRP: new insights into fragile X syndrome. Nat. Rev. Neurosci. 22, 209–222. 10.1038/s41583-021-00432-0.

16. Berry-Kravis, E.M., Lindemann, L., Jønch, A.E., Apostol, G., Bear, M.F., Carpenter, R.L., Crawley, J.N., Curie, A., Des Portes, V., Hossain, F., et al. (2018). Drug development for neurodevelopmental disorders: lessons learned from fragile X syndrome. Nat. Rev. Drug Discov. 17, 280–299. 10.1038/nrd.2017.221.

17. Protic, D., and Hagerman, R.J. (2024). State-of-the-art therapies for fragile X syndrome. Dev. Med. Child Neurol. 66, 863–871. 10.1111/dmcn.15885.

18. Laggerbauer, B., Ostareck, D., Keidel, E.M., Ostareck-Lederer, A., and Fischer, U. (2001). Evidence that fragile X mental retardation protein is a negative regulator of translation. Hum. Mol. Genet. 10, 329–338. 10.1093/hmg/10.4.329.

19. Li, Z., Zhang, Y., Ku, L., Wilkinson, K.D., Warren, S.T., and Feng, Y. (2001). The fragile X mental retardation protein inhibits translation via interacting with mRNA. Nucleic Acids Res. 29, 2276–2283. 10.1093/nar/29.11.2276.

20. Zhang, Y.Q., Bailey, A.M., Matthies, H.J., Renden, R.B., Smith, M.A., Speese, S.D., Rubin, G.M., and Broadie, K. (2001). Drosophila fragile X-related gene regulates the MAP1B homolog Futsch to control synaptic structure and function. Cell 107, 591–603. 10.1016/s0092-8674(01)00589-x.

21. Mazroui, R., Huot, M.E., Tremblay, S., Filion, C., Labelle, Y., and Khandjian, E.W. (2002). Trapping of messenger RNA by Fragile X Mental Retardation protein into cytoplasmic granules induces translation repression. Hum. Mol. Genet. 11, 3007–3017. 10.1093/hmg/11.24.3007.

22. Kurosaki, T., Cho, H., Abshire, E.T., Pröschel, C., Mitsutomi, S., Sato, H., Simko, E.A.J., Fraser, C.S., Sakano, H., and Maquat, L.E. (2025). FMRP drives mRNP targets into translationally silenced complexes. Mol. Cell 85, 2956–2972.e10. 10.1016/j.molcel.2025.06.012.

23. Das Sharma, S., Metz, J.B., Li, H., Hobson, B.D., Hornstein, N., Sulzer, D., Tang, G., and Sims, P.A. (2019). Widespread alterations in translation elongation in the brain of juvenile Fmr1 knockout mice. Cell Rep. 26, 3313–3322.e5. 10.1016/j.celrep.2019.02.086.

24. Coffee, R.L., Jr., Tessier, C.R., Woodruff, E.A., 3rd, and Broadie, K. (2010). Fragile X mental retardation protein has a unique, evolutionarily conserved neuronal function not shared with FXR1P or FXR2P. Dis. Model. Mech. 3, 471–485. 10.1242/dmm.004598.

25. Kiebler, M.A., and Bauer, K.E. (2024). RNA granules in flux: dynamics to balance physiology and pathology. Nat. Rev. Neurosci. 25, 711–725. 10.1038/s41583-024-00859-1.

26. Matheny, T., Rao, B.S., and Parker, R. (2019). Transcriptome-wide comparison of stress granules and P-bodies reveals that translation plays a major role in RNA partitioning. Mol. Cell. Biol. 39, e00313–19. 10.1128/MCB.00313-19.

27. Riggs, C.L., Kedersha, N., Ivanov, P., and Anderson, P. (2020). Mammalian stress granules and P bodies at a glance. J. Cell Sci. 133, jcs242487. 10.1242/jcs.242487.

28. Ayache, J., Bénard, M., Ernoult-Lange, M., Minshall, N., Standart, N., Kress, M., and Weil, D. (2015). P-body assembly requires DDX6 repression complexes rather than decay or Ataxin2/2L complexes. Mol. Biol. Cell 26, 2579–2595. 10.1091/mbc.E15-03-0136.

29. Minshall, N., Kress, M., Weil, D., and Standart, N. (2009). Role of p54 RNA helicase activity and its C-terminal domain in translational repression, P-body localization and assembly. Mol. Biol. Cell 20, 2464–2472. 10.1091/mbc.e09-01-0035.

30. Nakamura, A., Amikura, R., Hanyu, K., and Kobayashi, S. (2001). Me31B silences translation of oocyte-localizing RNAs through the formation of cytoplasmic RNP complex during Drosophila oogenesis. Development 128, 3233–3242. 10.1242/dev.128.17.3233.

31. Nakamura, A., Sato, K., and Hanyu-Nakamura, K. (2004). Drosophila Cup is an eIF4E binding protein that associates with Bruno and regulates oskar mRNA translation in oogenesis. Dev. Cell 6, 69–78. 10.1016/S1534-5807(03)00400-3.

32. Ripin, N., Macedo de Vasconcelos, L., Ugay, D.A., and Parker, R. (2024). DDX6 modulates P-body and stress granule assembly, composition, and docking. J. Cell Biol. 223, e202306022. 10.1083/jcb.202306022.

33. Wilhelm, J.E., Buszczak, M., and Sayles, S. (2005). Efficient protein trafficking requires trailer hitch, a component of a ribonucleoprotein complex localized to the ER in Drosophila. Dev. Cell 9, 675–685. 10.1016/j.devcel.2005.09.015.

34. Eulalio, A., Behm-Ansmant, I., Schweizer, D., and Izaurralde, E. (2007). P-body formation is a consequence, not the cause, of RNA-mediated gene silencing. Mol. Cell. Biol. 27, 3970–3981. 10.1128/MCB.00128-07.

35. Barbee, S.A., Estes, P.S., Cziko, A.-M., Hillebrand, J., Luedeman, R.A., Coller, J.M., Johnson, N., Howlett, I.C., Geng, C., Ueda, R., et al. (2006). Staufen- and FMRP-containing neuronal RNPs are structurally and functionally related to somatic P bodies. Neuron 52, 997–1009. 10.1016/j.neuron.2006.10.028.

36. Greenblatt, E.J., Obniski, R., Mical, C., and Spradling, A.C. (2019). Prolonged ovarian storage of mature Drosophila oocytes dramatically increases meiotic spindle instability. eLife 8, e49455. 10.7554/eLife.49455.

37. Hara, M., Lourido, S., Petrova, B., Lou, H.J., Von Stetina, J.R., Kashevsky, H., Turk, B.E., and Orr-Weaver, T.L. (2018). Identification of PNG kinase substrates uncovers interactions with the translational repressor TRAL in the oocyte-to-embryo transition. eLife 7, e33150. 10.7554/eLife.33150.

38. Kronja, I., Yuan, B., Eichhorn, S.W., Dzeyk, K., Krijgsveld, J., Bartel, D.P., and Orr-Weaver, T.L. (2014). Widespread changes in the posttranscriptional landscape at the Drosophila oocyte-to-embryo transition. Cell Rep. 7, 1495–1508. 10.1016/j.celrep.2014.05.002.

39. Voronina, E., Seydoux, G., Sassone-Corsi, P., and Nagamori, I. (2011). RNA granules in germ cells. Cold Spring Harb. Perspect. Biol. 3, a002774. 10.1101/cshperspect.a002774.

40. Weil, T.T., Parton, R.M., Herpers, B., Soetaert, J., Veenendaal, T., Xanthakis, D., Dobbie, I.M., Halstead, J.M., Hayashi, R., Rabouille, C., et al. (2012). Drosophila patterning is established by differential association of mRNAs with P bodies. Nat. Cell Biol. 14, 1305– 1313. 10.1038/ncb2627.

41. Protter, D.S.W., and Parker, R. (2016). Principles and properties of stress granules. Trends Cell Biol. 26, 668–679. 10.1016/j.tcb.2016.05.004.

42. Ivanov, P., Kedersha, N., and Anderson, P. (2019). Stress granules and processing bodies in translational control. Cold Spring Harb. Perspect. Biol. 11, a032813. 10.1101/cshperspect.a032813.

43. Smith, J., and Bartel, D.P. (2026). The G3BP stress-granule proteins reinforce the integrated stress response translation programme. Nat. Cell Biol. 28, 135–148. 10.1038/s41556-025-01834-3.

44. Glauninger, H., Wong Hickernell, C.J., Bard, J.A.M., and Drummond, D.A. (2022). Stressful steps: progress and challenges in understanding stress-induced mRNA condensation and accumulation in stress granules. Mol. Cell 82, 2544–2556. 10.1016/j.molcel.2022.05.014.

45. Putnam, A., Thomas, L., and Seydoux, G. (2023). RNA granules: functional compartments or incidental condensates? Genes Dev. 37, 354–376. 10.1101/gad.350518.123.

46. Götze, M., Dufourt, J., Ihling, C., Rammelt, C., Pierson, S., Sambrani, N., Temme, C., Sinz, A., Simonelig, M., and Wahle, E. (2017). Translational repression of the Drosophila nanos mRNA involves the RNA helicase Belle and RNA coating by Me31B and Trailer hitch. RNA 23, 1552–1568. 10.1261/rna.062208.117.

47. Khong, A., Matheny, T., Jain, S., Mitchell, S.F., Wheeler, J.R., and Parker, R. (2017). The stress granule transcriptome reveals principles of mRNA accumulation in stress granules. Mol. Cell 68, 808–820.e5. 10.1016/j.molcel.2017.10.015.

48. Hubstenberger, A., Courel, M., Bénard, M., Souquere, S., Ernoult-Lange, M., Chouaib, R., Yi, Z., Morlot, J.-B., Munier, A., Fradet, M., et al. (2017). P-body purification reveals the condensation of repressed mRNA regulons. Mol. Cell 68, 144–157.e5. 10.1016/j.molcel.2017.09.003.

49. Padrón, A., Iwasaki, S., and Ingolia, N.T. (2019). Proximity RNA labeling by APEX-seq reveals the organization of translation initiation complexes and repressive RNA granules. Mol. Cell 75, 875–887.e5. 10.1016/j.molcel.2019.07.030.

50. Hyman, S.L., Shores, A., and North, K.N. (2005). The nature and frequency of cognitive deficits in children with neurofibromatosis type 1. Neurology 65, 1037–1044. 10.1212/01.wnl.0000179303.72345.ce.

51. Kheradmand Kia, S., Verbeek, E., Engelen, E., Schot, R., Poot, R.A., de Coo, I.F.M., Lequin, M.H., Poulton, C.J., Pourfarzad, F., Grosveld, F.G., et al. (2012). RTTN mutations link primary cilia function to organization of the human cerebral cortex. Am. J. Hum. Genet. 91, 533–540. 10.1016/j.ajhg.2012.07.008.

52. Moortgat, S., Berland, S., Aukrust, I., Maystadt, I., Baker, L., Benoit, V., Caro-Llopis, A., Cooper, N.S., Debray, F.G., Faivre, L., et al. (2018). HUWE1 variants cause dominant X-linked intellectual disability: a clinical study of 21 patients. Eur. J. Hum. Genet. 26, 64–74. 10.1038/s41431-017-0038-6.

53. Zhang, J., Gambin, T., Yuan, B., Szafranski, P., Rosenfeld, J.A., Al Balwi, M., Alswaid, A., Al-Gazali, L., Al Shamsi, A.M., Komara, M., et al. (2017). Haploinsufficiency of the E3 ubiquitin-protein ligase gene TRIP12 causes intellectual disability with or without autism spectrum disorders, speech delay, and dysmorphic features. Hum. Genet. 136, 377–386. 10.1007/s00439-017-1763-1.

54. De Boulle, K., Verkerk, A.J.M.H., Reyniers, E., Vits, L., Hendrickx, J., Van Roy, B., Van den Bos, F., de Graaff, E., Oostra, B.A., and Willems, P.J. (1993). A point mutation in the FMR-1 gene associated with fragile X mental retardation. Nat. Genet. 3, 31–35. 10.1038/ng0193-31.

55. Myrick, L.K., Nakamoto-Kinoshita, M., Lindor, N.M., Kirmani, S., Cheng, X., and Warren, S.T. (2014). Fragile X syndrome due to a missense mutation. Eur. J. Hum. Genet. 22, 1185–1189. 10.1038/ejhg.2013.311.

56. Wan, L., Dockendorff, T.C., Jongens, T.A., and Dreyfuss, G. (2000). Characterization of dFMR1, a Drosophila melanogaster homolog of the fragile X mental retardation protein. Mol. Cell. Biol. 20, 8536–8547. 10.1128/MCB.20.22.8536-8547.2000.

57. Lee, F.C.Y., Chakrabarti, A.M., Hänel, H., Monzón-Casanova, E., Hallegger, M., Militti, C., Capraro, F., Sadée, C., Toolan-Kerr, P., Wilkins, O., et al. (2021). An improved iCLIP protocol. Preprint at bioRxiv. 10.1101/2021.08.27.457890.

58. Kurosaki, T., Mitsutomi, S., Hewko, A., Akimitsu, N., and Maquat, L.E. (2022). Integrative omics indicate FMRP sequesters mRNA from translation and deadenylation in human neuronal cells. Mol. Cell 82, 4564–4581.e11. 10.1016/j.molcel.2022.10.018.

59. Kara, E., McCambridge, A., Proffer, M., Dilts, C., Pumnea, B., Eshak, J., Smith, K.A., Fielder, I., Doyle, D.A., Ortega, B.M., et al. (2023). Mutational analysis of the functional motifs of the DEAD-box RNA helicase Me31B/DDX6 in Drosophila germline development. FEBS Lett. 597, 1848–1867. 10.1002/1873-3468.14668.

60. Liu, F., Putnam, A., and Jankowsky, E. (2008). ATP hydrolysis is required for DEAD-box protein recycling but not for duplex unwinding. Proc. Natl. Acad. Sci. USA 105, 20209–20214. 10.1073/pnas.0811115106.

61. Antar, L.N., Afroz, R., Dictenberg, J.B., Carroll, R.C., and Bassell, G.J. (2004). Metabotropic glutamate receptor activation regulates fragile X mental retardation protein and Fmr1 mRNA localization differentially in dendrites and at synapses. J. Neurosci. 24, 2648–2655. 10.1523/JNEUROSCI.0099-04.2004.

62. Rosario, R., Filis, P., Tessyman, V., Kinnell, H., Childs, A.J., Gray, N.K., and Anderson, R.A. (2016). FMRP associates with cytoplasmic granules at the onset of meiosis in the human oocyte. PLoS ONE 11, e0163987. 10.1371/journal.pone.0163987.

63. Fenton, A.R., Peng, R., Bond, C., Hugelier, S., Lakadamyali, M., Chang, Y.-W., Holzbaur, E.L.F., and Jongens, T.A. (2024). FMRP regulates MFF translation to locally direct mitochondrial fission in neurons. Nat. Cell Biol. 26, 2061–2074. 10.1038/s41556-024-01544-2.

64. Kang, J.-Y., Wen, Z., Pan, D., Zhang, Y., Li, Q., Zhong, A., Yu, X., Wu, Y.-C., Chen, Y., Zhang, X., et al. (2022). LLPS of FXR1 drives spermiogenesis by activating translation of stored mRNAs. Science 377, eabj6647. 10.1126/science.abj6647.

65. Mermod, J.J., and Crippa, M. (1978). Variations in the amount of polysomes in mature oocytes of Drosophila melanogaster. Dev. Biol. 66, 586–592. 10.1016/0012-1606(78)90264-6.

66. Grabarczyk, D.B., Ehrmann, J.F., Murphy, P., Yang, W.S., Kurzbauer, R., Bell, L.E., Deszcz, L., Neuhold, J., Schleiffer, A., Shulkina, A., et al. (2025). Architecture of the UBR4 complex, a giant E4 ligase central to eukaryotic protein quality control. Science 389, 909– 914. 10.1126/science.adv9309.

67. Tamanini, F., van Unen, L., Bakker, C., Sacchi, N., Galjaard, H., Oostra, B.A., and Hoogeveen, A.T. (1999). Oligomerization properties of fragile-X mental-retardation protein (FMRP) and the fragile-X-related proteins FXR1P and FXR2P. Biochem. J. 343, 517–523. PMID: 10527928

68. Youn, J.-Y., Dunham, W.H., Hong, S.J., Knight, J.D.R., Bashkurov, M., Chen, G.I., Bagci, H., Rathod, B., MacLeod, G., Eng, S.W.M., et al. (2018). High-density proximity mapping reveals the subcellular organization of mRNA-associated granules and bodies. Mol. Cell 69, 517–532.e11. 10.1016/j.molcel.2017.12.020.

69. Das, R., Schwintzer, L., Vinopal, S., Aguado Roca, E., Sylvester, M., Oprisoreanu, A.-M., Schoch, S., Bradke, F., and Broemer, M. (2019). New roles for the de-ubiquitylating enzyme OTUD4 in an RNA-protein network and RNA granules. J. Cell Sci. 132, jcs229252. 10.1242/jcs.229252.

70. Chang, C.-T., Bercovich, N., Loh, B., Jonas, S., and Izaurralde, E. (2014). The activation of the decapping enzyme DCP2 by DCP1 occurs on the EDC4 scaffold and involves a conserved loop in DCP1. Nucleic Acids Res. 42, 5217–5233. 10.1093/nar/gku129.

71. The ENCODE Project Consortium (2012). An integrated encyclopedia of DNA elements in the human genome. Nature 489, 57–74. 10.1038/nature11247.

72. Di Stefano, B., Luo, E.-C., Haggerty, C., Aigner, S., Charlton, J., Brumbaugh, J., Ji, F., Rabano Jiménez, I., Clowers, K.J., Huebner, A.J., et al. (2019). The RNA helicase DDX6 controls cellular plasticity by modulating P-body homeostasis. Cell Stem Cell 25, 622– 638.e13. 10.1016/j.stem.2019.08.018.

73. Luo, E.-C., Nathanson, J.L., Tan, F.E., Schwartz, J.L., Schmok, J.C., Shankar, A., Markmiller, S.J., Yee, B.A., Sathe, S., Pratt, G.A., et al. (2020). Large-scale tethered function assays identify factors that regulate mRNA stability and translation. Nat. Struct. Mol. Biol. 27, 989–1000. 10.1038/s41594-020-0477-6.

74. Van Nostrand, E.L., Freese, P., Pratt, G.A., Wang, X., Wei, X., Xiao, R., Blue, S.M., Chen, J.-Y., Cody, N.A.L., Dominguez, D., et al. (2020). A large-scale binding and functional map of human RNA-binding proteins. Nature 583, 711–719. 10.1038/s41586-020-2077-3.

75. Rhine, K., Li, R., Kopalle, H.M., Rothamel, K., Ge, X., Epstein, E., Mizrahi, O., Madrigal, A.A., Her, H.-L., Gomberg, T.A., et al. (2025). Neuronal aging causes mislocalization of splicing proteins and unchecked cellular stress. Nat. Neurosci. 28, 1174–1184. 10.1038/s41593-025-01952-z.

76. Rajyaguru, P., She, M., and Parker, R. (2012). Scd6 targets eIF4G to repress translation: RGG motif proteins as a class of eIF4G-binding proteins. Mol. Cell 45, 244–254. 10.1016/j.molcel.2011.11.026.

77. Zeidan, Q., He, F., Zhang, F., Zhang, H., Jacobson, A., and Hinnebusch, A.G. (2018). Conserved mRNA-granule component Scd6 targets Dhh1 to repress translation initiation and activates Dcp2-mediated mRNA decay in vivo. PLoS Genet. 14, e1007806. 10.1371/journal.pgen.1007806.

78. Baierlein, C., Hackmann, A., Gross, T., Henker, L., Hinz, F., and Krebber, H. (2013). Monosome formation during translation initiation requires the serine/arginine-rich protein Npl3. Mol. Cell. Biol. 33, 4811–4823. 10.1128/MCB.00873-13.

79. Ripin, N., and Parker, R. (2023). Formation, function, and pathology of RNP granules. Cell 186, 4737–4756. 10.1016/j.cell.2023.09.006.

80. Kedersha, N., Stoecklin, G., Ayodele, M., Yacono, P., Lykke-Andersen, J., Fritzler, M.J., Scheuner, D., Kaufman, R.J., Golan, D.E., and Anderson, P. (2005). Stress granules and processing bodies are dynamically linked sites of mRNP remodeling. J. Cell Biol. 169, 871–884. 10.1083/jcb.200502088.

81. Kedersha, N., Panas, M.D., Achorn, C.A., Lyons, S., Tisdale, S., Hickman, T., Thomas, M., Lieberman, J., McInerney, G.M., Ivanov, P., et al. (2016). G3BP–Caprin1–USP10 complexes mediate stress granule condensation and associate with 40S subunits. J. Cell Biol. 212, 845–860. 10.1083/jcb.201508028.

82. Mat□j□, D., Eichenberger, B., Voigt, F., Eglinger, J., Roth, G., and Chao, J.A. (2020). Single-molecule imaging reveals translation of mRNAs localized to stress granules. Cell 183, 1801–1812.e13. 10.1016/j.cell.2020.11.010.

83. Laver, J.D., Ly, J., Winn, J.K., Karaiskakis, A., Lin, S., Nie, K., Benic, G., Jaberi-Lashkari, N., Cao, W.X., Khademi, A., et al. (2020). The RNA-binding protein Rasputin/G3BP enhances the stability and translation of its target mRNAs. Cell Rep. 30, 3353–3367.e7. 10.1016/j.celrep.2020.02.066.

84. Zaessinger, S., Busseau, I., and Simonelig, M. (2006). Oskar allows nanos mRNA translation in Drosophila embryos by preventing its deadenylation by Smaug/CCR4. Development 133, 4573–4583. 10.1242/dev.02649.

85. Chen, R., Stainier, W., Dufourt, J., Lagha, M., and Lehmann, R. (2024). Direct observation of translational activation by a ribonucleoprotein granule. Nat. Cell Biol. 26, 1322–1335. 10.1038/s41556-024-01452-5.

86. McCoy, M.J., and Fire, A.Z. (2020). Intron and gene size expansion during nervous system evolution. BMC Genomics 21, 360. 10.1186/s12864-020-6760-4.

87. McCoy, M.J., and Fire, A.Z. (2024). Parallel gene size and isoform expansion of ancient neuronal genes. Curr. Biol. 34, 1635–1645. 10.1016/j.cub.2024.02.021.

88. Bassell, G.J., and Warren, S.T. (2008). Fragile X syndrome: loss of local mRNA regulation alters synaptic development and function. Neuron 60, 201–214. 10.1016/j.neuron.2008.10.004.

89. Giles, A.C., and Grill, B. (2020). Roles of the HUWE1 ubiquitin ligase in nervous system development, function and disease. Neural Dev. 15, 6. 10.1186/s13064-020-00143-9.

90. Barnsby-Greer, L., Mabbitt, P.D., Dery, M.-A., Squair, D.R., Wood, N.T., Lamoliatte, F., Lange, S.M., and Virdee, S. (2024). UBE2A and UBE2B are recruited by an atypical E3 ligase module in UBR4. Nat. Struct. Mol. Biol. 31, 351–363. 10.1038/s41594-023-01192-4.

91. Báez-Flores, J., Rodríguez-Martín, M., and Lacal, J. (2023). The therapeutic potential of neurofibromin signaling pathways and binding partners. Commun. Biol. 6, 436. 10.1038/s42003-023-04815-0.

92. Shevchenko, A., Wilm, M., Vorm, O., and Mann, M. (1996). Mass spectrometric sequencing of proteins from silver-stained polyacrylamide gels. Anal. Chem. 68, 850–858. 10.1021/ac950914h.

93. Maurin, T., Lebrigand, K., Castagnola, S., Paquet, A., Jarjat, M., Popa, A., Grossi, M., Rage, F., and Bardoni, B. (2018). HITS-CLIP in various brain areas reveals new targets and new modalities of RNA binding by fragile X mental retardation protein. Nucleic Acids Res. 46, 6344–6355. 10.1093/nar/gky267.

94. Su, S., Xue, Y., Sharov, A., Zhang, Y., Lee, S.K., Martindale, J.L., Li, W., Ku, W.L., Zhao, K., De, S., et al. (2022). A dual-activity topoisomerase complex regulates mRNA translation and turnover. Nucleic Acids Res. 50, 7013–7033. 10.1093/nar/gkac538.

95. Teo, G., Liu, G., Zhang, J., Nesvizhskii, A.I., Gingras, A.-C., and Choi, H. (2014). SAINTexpress: improvements and additional features in Significance Analysis of INTeractome software. J. Proteomics 100, 37–43. 10.1016/j.jprot.2013.10.023.

96. Hornstein, N., Torres, D., Das Sharma, S., Tang, G., Canoll, P., and Sims, P.A. (2016). Ligation-free ribosome profiling of cell type-specific translation in the brain. Genome Biol. 17, 149. 10.1186/s13059-016-1005-1.

97. Dunn, J.G., Foo, C.K., Belletier, N.G., Gavis, E.R., and Weissman, J.S. (2013). Ribosome profiling reveals pervasive and regulated stop codon readthrough in Drosophila melanogaster. eLife 2, e01179. 10.7554/eLife.01179.

98. Love, M.I., Huber, W., and Anders, S. (2014). Moderated estimation of fold change and dispersion for RNA-seq data with DESeq2. Genome Biol. 15, 550. 10.1186/s13059-014-0550-8.

